# Defining the contribution of *Troy*-positive progenitor cells to the mouse esophageal epithelium

**DOI:** 10.1101/2024.01.11.575258

**Authors:** David Grommisch, Menghan Wang, Evelien Eenjes, Maja Svetličič, Qiaolin Deng, Pontus Giselsson, Maria Genander

**Affiliations:** Department of Cell and Molecular Biology, Karolinska Institutet, Sweden; Department of Physiology and Pharmacology, Karolinska Institutet, Sweden; Department of Automatic Control, Lund University, Sweden

## Abstract

Progenitor cells adapt their behavior in response to tissue demands. However, the molecular mechanisms controlling esophageal progenitor decisions remain largely unknown. Here we demonstrate the presence of a *Troy* (*Tnfrsf19*)-expressing progenitor subpopulation localized to defined regions along the mouse esophageal axis. Lineage tracing and mathematical modelling demonstrate that *Troy-*positive progenitor cells are prone to undergoing symmetrical fate choices and contribute to esophageal tissue homeostasis long-term. Functionally, TROY inhibits progenitor proliferation and enables commitment to differentiation without affecting fate symmetry. Whereas *Troy* expression is stable during esophageal homeostasis, progenitor cells downregulate *Troy* in response to tissue stress, enabling proliferative expansion of basal cells refractory to differentiation and reestablishment of tissue homeostasis. Our results demonstrate functional, spatially restricted, progenitor heterogeneity in the esophageal epithelium and identify how dynamic regulation of *Troy* coordinates tissue generation.

## Introduction

While many epithelial tissues are maintained by long-lasting stem cells, the stratified squamous epithelium of the esophagus is maintained by proliferation of relatively short-lived progenitor cells residing in the basal layer, which upon commitment to differentiation exit the cell cycle, stratify from the basal layer and eventually shed off at the tissue surface.

Early work characterizing basal cells in the esophagus describes functionally distinct cell states when progenitor subpopulations are challenged *in vitro* ^1-4^. In contrast, *in vivo* lineage tracing of sparsely labelled basal cells followed by mathematical modelling conclude that the esophageal epithelium, much like the epidermis and oral epithelium, is sustained by a single, functionally equivalent progenitor population ^5-7^, maintaining stratified epithelial homeostasis through stochastic fate decisions. Each progenitor cell division has thus the potential to generate daughter cells of either symmetric fate (progenitor-progenitor or differentiated-differentiated cell) or two daughter cells with distinct fate (progenitor-differentiated cell), symmetric fate being the less likely outcome ^5-7^.

Although the single-progenitor model is attractive in simplicity and robustness as well as able to mechanistically explain esophageal tissue homeostasis at large, the random targeting strategy employed does not formally exclude the possibility that subpopulations of progenitor cells exist. Strategies directed to label specific epidermal cell populations have unearthed cells with fate bias ^8,9^ and it is well established that subsets of skin and intestinal progenitor cells act to diversify the tissue response upon regenerative or oncogenic cues ^10,11^. Lack of robust data has so far precluded similar detailing in the esophageal epithelium.

Esophageal basal cells eventually undergo a coordinated program of differentiation in order to maintain tissue integrity ^2,12-15^. Differentiation is largely mediated by Notch and BMP signaling pathways ^2,12,13,15^ acting to promote cell cycle exit and induce stratification of individual cells during homeostasis. Tissue-wide biomechanical forces act upstream of signaling effectors to fine-tune commitment to differentiation during esophageal development^14^ and adult esophageal progenitor cells respond to tissue stress by transiently altering cell division and stratification rates ^7^, indicating that balancing proliferation and differentiation is a general mechanism in place to accommodate increased need for esophageal tissue production. Identification of molecular effectors allowing equivalently fated cells to independently and dynamically balance proliferation and differentiation are however largely lacking.

*Troy (Tnfrsf19)* delineates stem and progenitor cells in multiple tissues ^16-21^. Whereas *Troy* demarks rapidly proliferating intestinal stem and epidermal progenitor cells, *Troy* is also found in quiescent ependymal cells in the spinal cord and in differentiated chief cells in the gastric corpus, which both retain the ability to proliferate in response to injury and contribute to tissue regeneration ^18,20^. Expression of *Troy* is thus not associated with a specific stem or progenitor state, and it seems likely that the function of TROY is context dependent, reflecting cell type specific interaction with additional receptors ^16,22,23^ or activation of distinct downstream signaling pathways ^24,25^.

Here, we identify *Troy* as a marker for differentially fated progenitor cells and a negative effector of progenitor proliferation, acting as a molecular sensor to accommodate tissue homeostasis along the esophageal axis.

## Results

### *Troy* delineates a regionally restricted population of basal cells

Quantitative studies have suggested that the esophagus is maintained by a single functionally equivalent progenitor population ^6,7^. A single dose of tamoxifen to *Troy-CreER^T2^:tdTomato* (Troy^WT/KI^) mice resulted in tdTOMATO reporter expression in basal cells distributed in a proximal-to-distal gradient along the esophageal axis (Figure 1A-B). Localization of *Troy* mRNA mirrored the unequal distribution of tdTOMATO reporter-positive basal cells (Figure 1C and S1A), suggesting that the reporter expression faithfully recapitulates endogenous *Troy* expression. In addition to the proximal-to-distal *Troy-*gradient, tdTOMATO expression and *Troy* mRNA was found to be unequally distributed around the esophageal circumference, where regions of high tdTOMATO reporter or mRNA expression commonly localized to epithelium pointed towards the lumen of the esophageal tube (Figure 1D-E and S1A). A subset of *Pdgfr*α:H2BeGFP-positive stromal fibroblasts also expressed tdTOMATO (Figure S1B).

**Figure 1.**
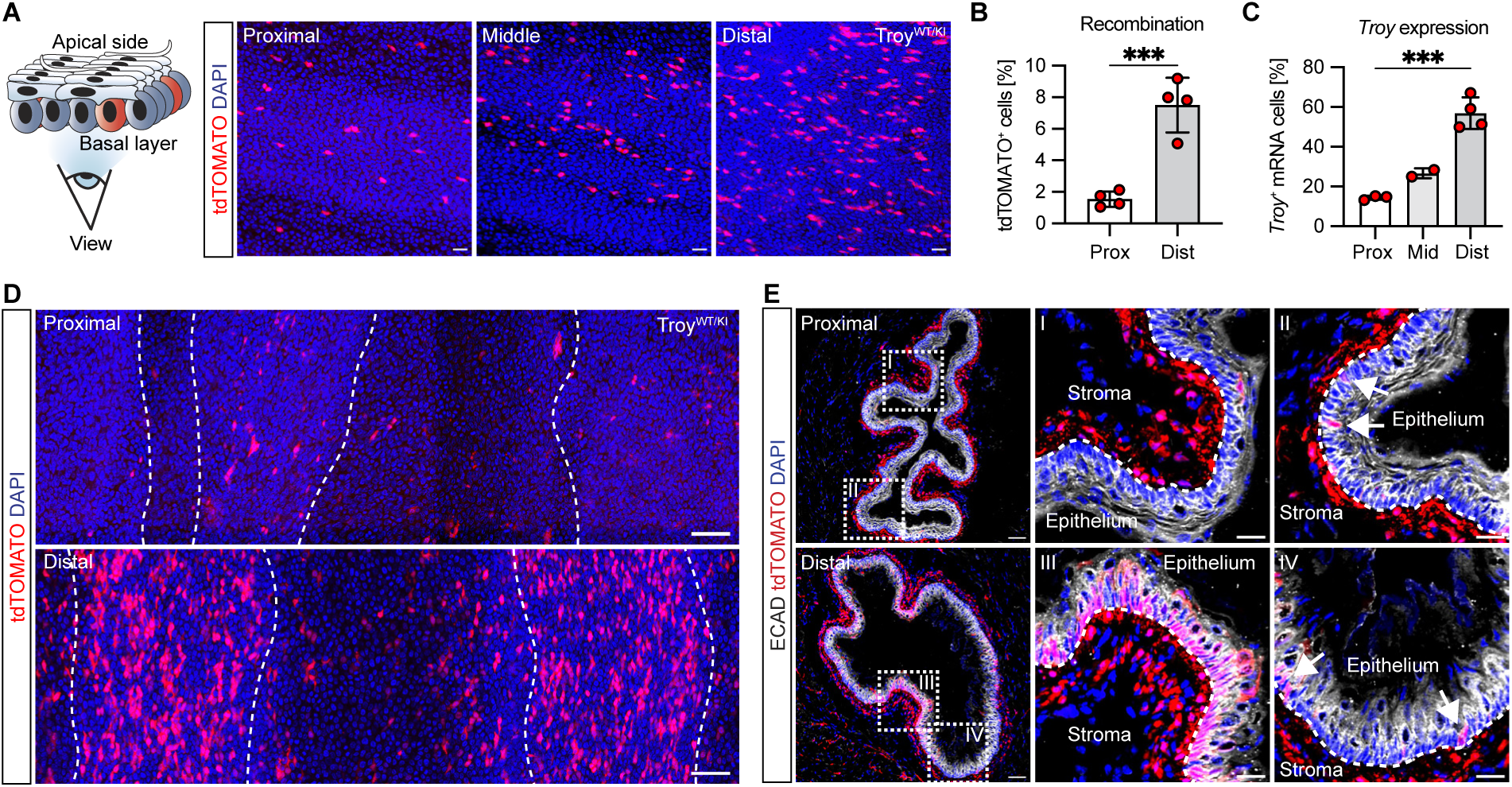
*Troy* delineates a regionally restricted population of basal cells. **(A)** Distribution of *Troy-CreER^T2^:tdTomato* (Troy^WT/KI^) reporter-positive cells along the esophageal axis. **(B)** Percentage of tdTOMATO positive basal cells in proximal and distal esophageal whole mount preparations. **(C)** Quantification of *Troy* mRNA-positive cells in cross sections. **(D-E)** *Troy* reporter (tdTOMATO) positive cells are enriched in specific distal epithelial regions. White lines **(D)** delineate the folding of the epithelium as seen in whole mount preparations. **(E)** Cross sections from proximal and distal esophagus revealing the spatial enrichment of tdTOMATO reporter expression in specific regions. Data are represented as mean ± SD. ^∗∗∗^p < 0.001 Statistical tests are: Two-sided unpaired t-tests (**B** and **C**). Scale bars are: **A** and inlets 20 μm, (**D**) 100 μm and (**E**) 50 μm.

Considering the local enrichment of *Troy-*expressing basal cells, we wondered if reporter expression reflects a specific, and perhaps transient, basal cell state. To this end, we administered tamoxifen once weekly for three consecutive weeks and assessed recombination efficiency and clone size (S1C-H). We found that regions of alternating reporter expression were maintained (Figure S1F) and the percentage of recombined cells as well as the average clone sizes were unaltered with repetitive tamoxifen administration (Figure S1D-E and S1G-H), indicating that repetitive injections targeted predominantly the same subpopulation of basal cells at all three timepoints. To understand if *Troy* expression correlated to a specific cell cycle phase, we administered tamoxifen to Troy^WT/KI^ mice at night, a time point when esophageal basal cell proliferation is low due to the circadian rhythm ^26^. However, the percentage of tdTOMATO reporter-positive, or *Troy* expressing, cells was similar when comparing Troy^WT/KI^ recombination at night or day (Figure S1I-L). In addition, we combined a 2-hour EdU pulse (S-phase) with CYCLINA2 immunofluorescence (S and G2 expression), allowing us to identify CYCLINA2-positive, EdU-negative progenitor cells in G2-phase (Figure S1M-N). We found that *Troy*-positive and negative basal cells were equally likely to be in G2. Collectively, these data argue against a cell cycle stage restricted expression of *Troy* and suggest that *Troy* expression delineates a distinct subset of esophageal basal cells localized to discrete epithelial regions.

### *Troy* marks progenitor cells with a distinct transcriptional signature

To characterize the nature of *Troy*-expressing basal cells, we turned to 10X Genomics single-cell sequencing. In contrast to previous published single-cell data sets ^14,27^, we enriched for progenitor cells by FACS isolating basal (CD104 (*Itgb4*) and CD49f (*Itga6*) double positive) cells, using wild type (Troy^WT/WT^) mice. RaceID ^28^ identified 7 clusters representing discrete stages of the esophageal progenitor cell differentiation trajectory, in line with previous reports ^27^ (Figure 2A-B, S2A), (Supplemental Table 1). Interestingly, *Troy-*positive cells did not cluster separately, but were enriched in *Krt5*/*Trp63* basal clusters 1 and 6 (9.3% and 4.6% of basal cells respectively), where cluster 6 represented cycling (*Mki67*/*Cdk1*/*Top2a*) basal cells (Figure 2B-C and S2B-C, Supplemental Table 2). Additionally, some *Troy-*expressing cells were localized to cluster 3 and 2 (4.1 and 1.03% of cells respectively), likely representing basal cells (*Krt5*/*Krt15*) committed to differentiation (*Krt4*, *Krtdap* and *Klf4*). In the remaining clusters (4, 7 and 5), the frequency of *Troy-*expressing cells was low and differentiation marker expression abundant (Figure 2B-C). Immunostaining confirmed that tdTOMATO-reporter positive and negative basal cells expressed comparable levels of established progenitor marker p63, KRT5 and SOX2 (Figure 2D and S2D), indicating that *Troy* expression is associated with an undifferentiated basal progenitor cell state.

**Figure 2.**
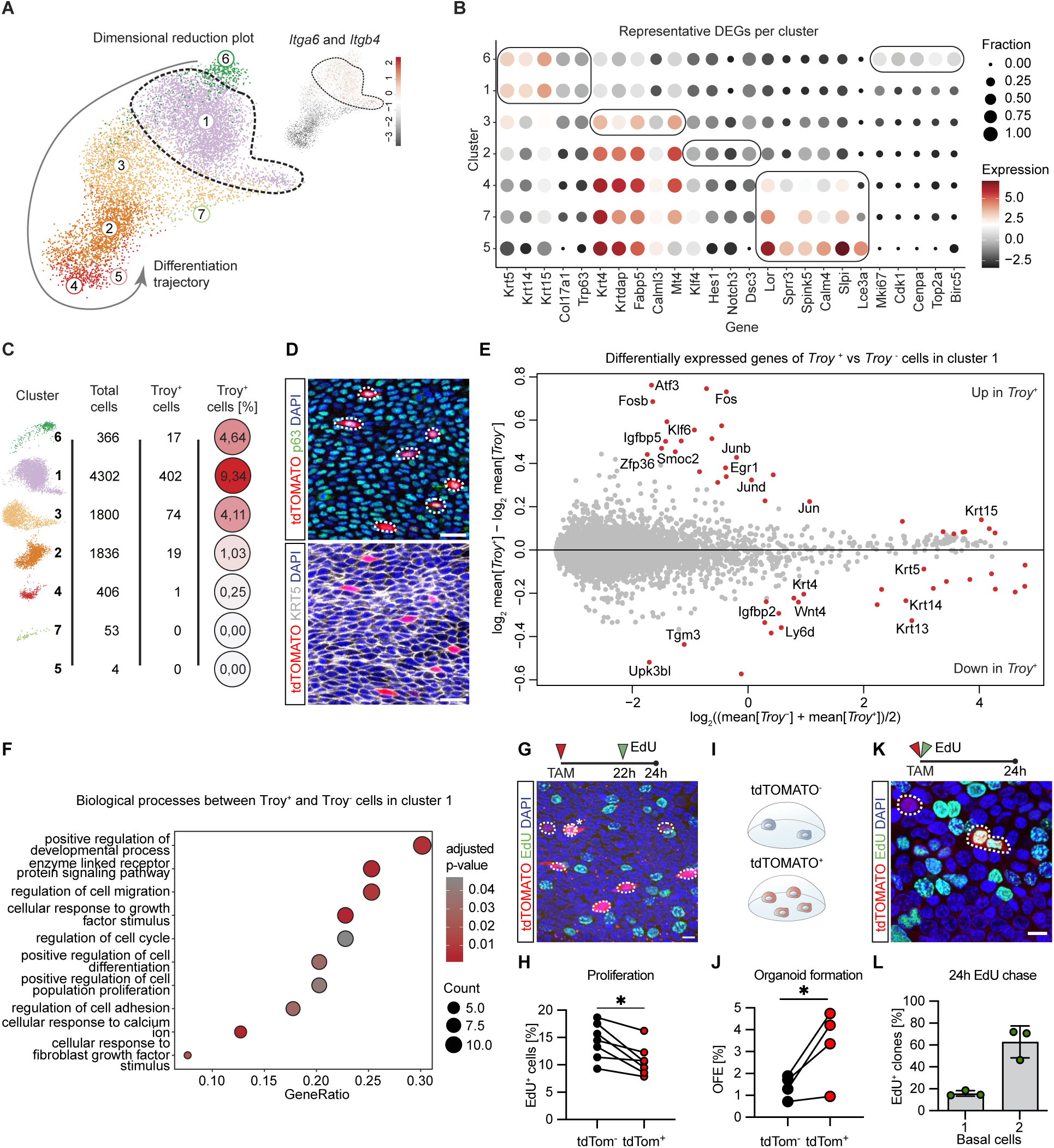
*Troy* marks a functionally distinct progenitor population. **(A)** Dimensional reduction plot of esophageal epithelial cells. Upper right corner highlights *Itga6* and *Itgb4* expressing progenitor cells **(B)** Representative differentially expressed genes for each cluster. **(C)** Most *Troy*-positive cells are found in undifferentiated basal cells (cluster 6, 1 and 3). **(D)** *Troy* reporter-positive cells in the basal layer express p63 and KRT5. **(E)** Volcano plot visualization of 54 differentially expressed genes comparing *Troy*-positive and negative progenitor cells in cluster 1. **(F)** Gene Ontology analysis (Biological process) based on differentially expressed genes in *Troy*-positive and negative cells in cluster 1. **(G-H)** *In vivo* administration of EdU for 2 hours. Pair-wise comparison within animals (n=7). **(I-J)** tdTOMATO-positive progenitors are better at forming organoids compared to tdTOMATO-negative progenitors. Organoid forming efficiency is compensated for sorting efficiency. **(K-L)** Joint administration of EdU and tamoxifen 24 hours before sacrifice identifies two-cell clones in the basal layer. Data are represented as mean ± SD. ^∗^p< 0.05, ^∗∗^p<0.01^∗∗∗^p < 0.001 Statistical tests are: Wilcoxon matched-pairs signed ranked test (**H**), Two-sided paired t-test (**J**). Scale bars are: (**H** and **K**) 10μm; (**D**) 20μm. n=4-7 in all experiments.

Focusing on progenitor cells enriched in cluster 1, we identified 54 genes differentially expressed in *Troy-*positive and negative progenitor cells (Figure 2E)(Supplemental Table 3). Gene ontology analysis revealed that differentially expressed genes associated with core cellular processes such as regulation of proliferation and differentiation and response to growth factors (Figure 2F and S2E)(Supplemental Table 4). Probing deeper into the 54 differentially expressed genes, we found a general enrichment of transcriptional regulators associated with immediate-early response (*Jun, Junb, Jund, Fos, Fosb, Atf3* and *Egr1*) indicating that *Troy-* positive progenitors engage distinct transcriptional programs. We also identified downregulation of several genes associated with epithelial differentiation (*Krt13, Krt4, Tgm3, Upk3bl* and *Ly6d*), suggesting that *Troy*-positive cells do not represent a subset of progenitor cells committed to differentiation ^8,9^. Finally, we identified a group of genes (*Wnt4, Igfbp2, Igfbp5* and *Smoc2*) shown to be WNT targets, effectors or modulators ^29-31^, suggesting that WNT signaling could be differentially sensed by *Troy-*positive versus negative progenitor cells. Taken together, unbiased single-cell sequencing confirms *Troy* expression in basal progenitor cells, reveals that the global transcriptomes of *Troy*-positive and negative progenitor cells are close but identifies a repertoire of transcriptional regulators as *Troy-*progenitor signature genes.

### *Troy-*positive progenitors are functionally distinct

The single-cell data analysis suggested that proliferation could be differentially regulated in *Troy*-positive and -negative progenitor cells (Figure 2F and S2E and S2H). To understand if proliferation was altered in *Troy*-positive, compared to *Troy*-negative progenitor cells, we administered EdU for 2 hours *in vivo*. Quantification revealed that *Troy-*positive progenitors were less proliferative than their *Troy*-negative neighboring progenitor cells when performing pair-wise comparisons of the two populations within the same esophagus (Figure 2G-H). We then FACS isolated tdTOMATO-positive and negative basal cells 24 hours after tamoxifen induction and challenged both cell populations to form organoids (Figure S2F-G) ^2,32^. Isolated tdTOMATO*-*positive progenitor cells were enriched for *Troy* mRNA and displayed increased organoid forming efficiency when compared to tdTOMATO-negative progenitors (Figure 2I-J and S2I-K). A majority of adult esophageal progenitor cells divide along the basement membrane before committing to differentiation ^7,14^. To understand if *Troy*-positive progenitor cells follow the same behavior, we concomitantly administered EdU and tamoxifen. After 24 hours, over 60% of reporter-positive two-cell basal clones were also EdU-positive, indicating that they were derived from a single labelled progenitor cell that had divided parallel to the basement membrane (Figure 2K-L and S2L-M).

Our data suggest that Troy-progenitor cells are functionally distinct, both in terms of proliferation (Figure 2G-H) and renewal capacity (Figure 2J). To understand if the distal enrichment of *Troy*-positive progenitor cells correlates to regional differences in tissue dynamics, we compared the proximal and distal progenitor cell behavior in wild type mice. Labelling of proliferating cells using a 1-hour EdU pulse revealed no statistically significant difference in proliferation (p=0.089, n=5) when comparing the proximal and distal esophagus of the same mouse (Figure S2N). However, mapping cell fates by quantifying the ratio of all EdU-positive basal and suprabasal cells after 48 hours of tracing ^33^ revealed that distal EdU-labelled cells were more often found in the basal layer when compared to the proximal esophagus (Figure S2O). We then turned to analyzing clones composed of either only basal cells or only suprabasal cells as a proxy for fate choice. Quantification revealed no significant difference in clones consisting of only basal cells, whereas the fraction of suprabasal clones (with no basal contribution) was reduced in the distal compared to the proximal esophagus (Figure S2P-R). These observations demonstrate that local tissue dynamics contribute to maintenance of esophageal homeostasis.

### *Troy-*positive progenitors are more likely to generate symmetrically fated progeny

To probe the long-term tissue contribution of *Troy-*positive progenitor cells, we treated Troy^WT/KI^ mice with tamoxifen for three days and performed lineage tracing for up to one year (4 days followed by 1, 3, 6 and 12 months) (Figure 3A and S3A-B). Quantifying surviving *Troy-*derived clones in the upper esophagus, we found that the average basal clone size increased with time, reaching an average of around 40 basal cells after one year (Figure 3B). In line with described dynamics of neutral drift ^6,7^, the number of persisting basal clones in the epithelium decreased with time (Figure 3C) rendering the overall tissue contribution of labelled basal cells constant (Figure 3D). These data indicate that during esophageal homeostasis, *Troy-* positive and -negative progenitor cells compete neutrally.

**Figure 3.**
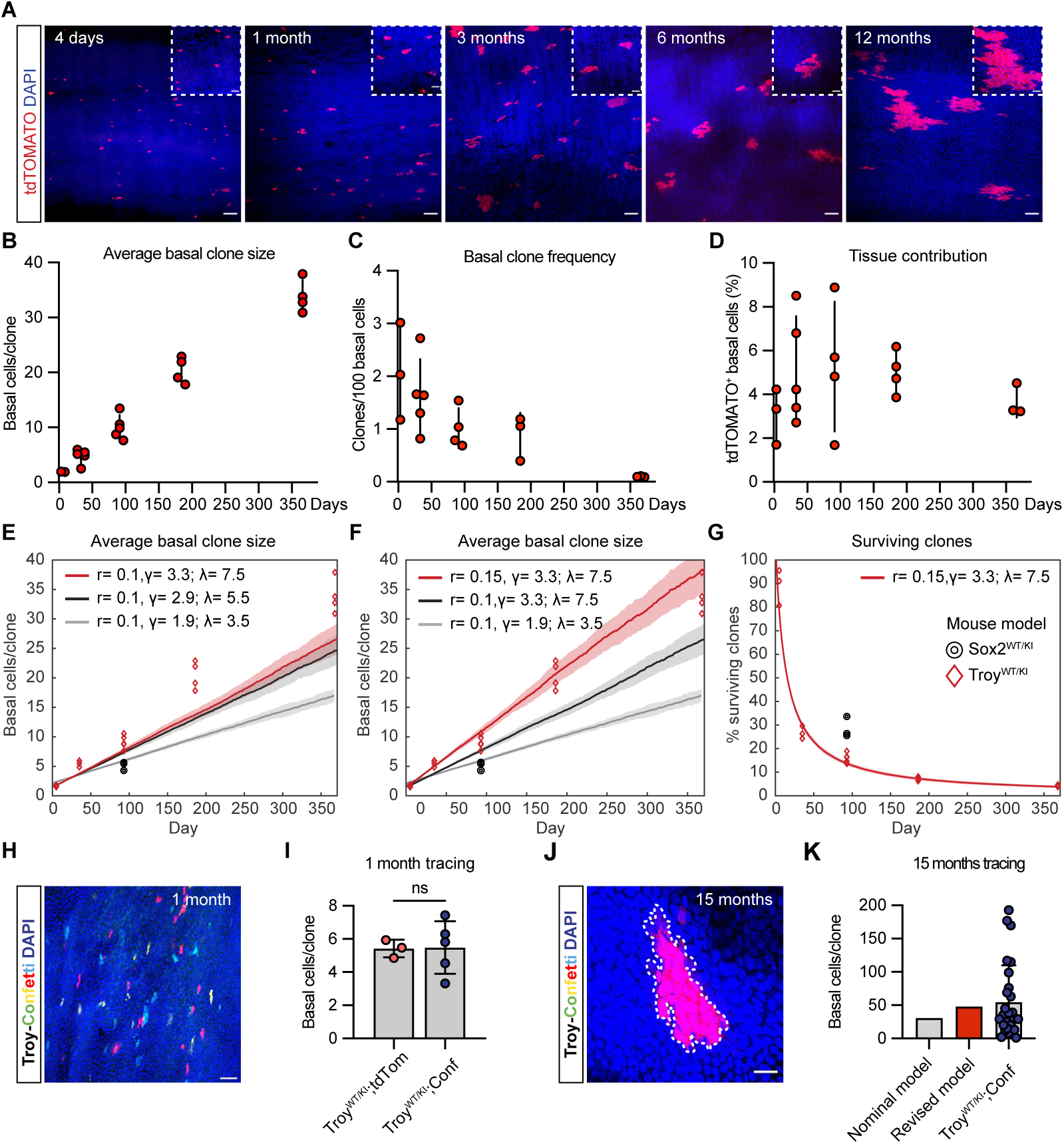
*Troy*-positive progenitors contribute to tissue homeostasis long-term. **(A)** Lineage tracing for 4 days, 1, 3, 6 and 12 months in *Troy-CreER^T2^:tdTomato* mice. **(B)** Quantification of average basal clone sizes. Each circle represents the average clone size in one animal. **(C)** Number of clones, as calculated by the clone number per 100 basal cells, decrease with time. **(D)** Tissue contribution of tdTOMATO-labelled clones over time is constant. **(E)** Basal clone sizes derived from *Troy-*positive progenitors (red diamonds) or Sox2^WT/KI^-tracing (black circles) compared to the single-progenitor model predictions: (grey line, *r*=0.1, λ=1.9, γ=3.5)^7^ and (black line *r*=0.1, λ=2.9, γ=5.5)^6^. The red line represents the predicted basal clone sizes using Troy^WT/KI^ determined parameters (*r*=0.1, λ=3.3, γ=7.5) based on the most conservative relative comparison ^6^. **(F)** Modeling of basal clone size comparing the nominal single-progenitor model (grey line, *r*=0.1, λ=1.9, γ=3.5)^7^ with Troy^WT/KI^ strain specific parameters (λ=3.3, γ=7.5) using *r*=0.1 (black line) and *r*=0.15 (red line). **(G)** Clones derived from *Troy-*positive cells (red diamonds) are lost faster than Sox2^WT/KI^-traced clones (black circles). Increasing fate probabilities (*r*=0.15, red line) explain the increased loss of clones. **(H)** Representative image of clone distribution in the *Troy-CreER^T2^:Confetti* reporter line 1 month after recombination. **(I)** Quantification of *Troy-CreER^T2^:Confetti* distal basal clone sized compared to upper *Troy-CreER^T2^:tdTomato* basal clone sizes. **(J-K)** Representative image and quantification of a single-colored clone at 15 months of tracing using the *Troy-CreER^T2^:Confetti* reporter line. Confetti clones are compared to the nominal (*r*=0.1, λ=2.9, γ=5.5) or revised (*r*=0.15, λ=3.3, γ=7.5) model parameters. Data are represented as mean ± SD. ns p > 0.05. Statistical tests are: Tow-sided unpaired t-test (**I**). Shaded areas in (**J** and **K**), indicate SD based on simulations. Scale bars are: (**A and H**) 50μm, (**A,** inlets) 20μm, (**J**) 20μm. n=4-7 in all experiments.

We compared our data with the nominal, single-progenitor model where the probability of equal fate choice (*r,* progenitor-progenitor or differentiated-differentiated fate) determines the likelihood for unequal (progenitor-differentiated) fate (1-2*r*). The nominal model is based on random labelling of basal cells, where a low probability for symmetrical fate choice (*r* **<**0.1) best explains the behavior of basal cells with experimentally determined weekly division (λ) and stratification (γ) rates ^6,7^.

Although the observed clone behavior fits the described stochastic model well, we found that the distribution of basal clone sizes was skewed towards larger basal clones compared to the nominal model prediction, using previously defined parameters (Figure S3C-F (λ=1.9 : γ=3.5), S3G-J (λ=2.9 : γ=5.5) and 3E, grey and black line) ^6,7^. To address if the observed differences in basal clone sizes could be explained by an overall altered esophageal tissue dynamic specific for the Troy^WT/KI^ mouse line, we first performed unbiased, random, labelling and lineage tracing of basal cells using the Sox2^WT/KI^ (*Sox2-CreER^T2^:YFP*) line. Random YFP reporter labelling of progenitor cells using Sox2^WT/KI^ would not only mimic the experimental rationale of published work ^6,7^, but also, due to the low abundance of *Troy-* positive progenitors in the upper esophagus, target largely *Troy-*negative progenitor cells. Administration of tamoxifen recombined cells uniformly over the esophageal axis (Figure S3K). Quantification of basal clone sizes at 3 months of tracing revealed that *Sox2*-derived basal clone sizes matched the model predictions (Figure 3E, black circles, grey and black lines), confirming the validity of the published model in describing expected basal clone sizes using an unbiased labelling method.

Leveraging the most conservative parameters of the single-progenitor model ^6,7^ (predicting the largest basal clone sizes)(Figure 3E, black line), we then performed an unbiased parameter scan to identify the optimal proliferation rate (λ) and fate choice probability (*r*) upon a fixed stratification rate (γ=5.5, nominal model) using the *Sox2*-derived average basal clone sizes. We identify *r* <0.1 to be the optimal fate probability within the suggested proliferative range (λ=2.9, nominal model) (Figure S3Q, white dashed lines), thereby recapitulating previous work using an unbiased labelling strategy ^6,7^.

To explore if mouse line specific differences in proliferation (λ) and stratification (γ) could explain the increased basal clone sizes observed when targeting *Troy*-positive progenitor cells, we compared proliferation and stratification dynamics in the Sox2^WT/KI^ and Troy^WT/KI^ mouse line by quantifying KI67, EdU and CYCLINA2 expression (Figure S3L-O) as well as the ratio of basal-to-suprabasal EdU-positive cells after 24 hours of chase (Figure S3P). We demonstrated that although the overall tissue dynamics of Troy^WT/KI^ is distinct compared to Sox2^WT/KI^ mouse line, higher proliferation is counteracted by increased stratification allowing for maintenance of tissue homeostasis.

Using our experimental data (Figure S3L-P), we then calculated the relative proliferation and stratification rates in Troy^WT/KI^ compared to Sox2^WT/KI^ mice (Figure S3L-P). Modelling basal clone sizes implementing the estimated Troy^WT/KI^ mouse line specific parameters (λ=3.3, γ=7.5) however failed to explain the observed Troy^WT/KI^ basal clone sizes (Figure 3E, red line), indicating that strain-specific differences in tissue dynamics alone are not enough to explain the observed clone behavior in Troy^WT/KI^ and suggesting that fate choice (*r*) could be altered.

Revisiting the unbiased parameter scan using average basal clone sizes derived from *Troy*-positive progenitor cells, using either γ=5.5 (nominal model) or γ=7.5 (Troy^WT/KI^ specific) at 3 months or combining all traced time points suggests that an increase in symmetric fate choice probability (*r* >0.1) drive increased basal clone sizes in the Troy^WT/KI^ line (Figure S3R-U, optimal *r* indicated with white dashed lines). Predicting the average basal clone sizes when increasing symmetric fate probabilities match *Troy*-derived clone sizes (Figure 3F and S3V) and demonstrate that symmetric fate probabilities in *Troy*-positive progenitors are likely higher than previously reported for esophageal progenitor cells.

Increasing the probability to generate symmetrical fates in combination with the constant tissue contribution observed from labelled *Troy*-derived cells (Figure 3D) infer that loss of basal clones through delamination must be increased. To this end, we found that clones emanating from *Troy*-positive progenitors are more likely to be competed out, and that the rate of clone loss can be explained by increased symmetric fate choice probabilities (Figure 3G). Collectively, these data suggest that *Troy-*positive progenitor cells are differentially fated in comparison to *Troy-*negative progenitor cells.

### *Troy*-derived clone sizes are not significantly skewed by clone fusion

To delineate if the observed large clone sizes could be explained by fusion of two neighboring smaller clones during tracing we exploited the increased resolution of the *Troy-CreER^T2^:Confetti* line, allowing for recombination in four colors. Because of the low recombination efficiency of the Confetti allele, we preferentially mark *Troy*-positive cells in the distal esophagus. This allowed us to compare the sizes of tdTOMATO-positive (*Troy-CreER^T2^:tdTomato)* basal clones in the upper to Confetti reporter-positive *(Troy-CreER^T2^:Confetti)* basal clones in the distal esophagus, after 1 month of tracing. All Confetti-clones included in the analysis were uniformly labelled by a single color and were not significantly different in their average clone size compared to the tdTOMATO-labelled basal clones in the upper esophagus (Figure 3H-I). After 15 months of tracing on the Confetti background, only a few, single-colored, clones remained (Figure 3J). Importantly, the basal clone sizes observed were larger than what would be predicted by the nominal model (Figure 3K). These data indicate that increased probability of symmetrical fate choice in *Troy*-positive progenitor cells is maintained over the esophageal axis.

### TROY is dispensable for symmetric fate choices

In an attempt to dissociate TROY-mediated effects on progenitor cell proliferation and fate choice, we generated *Troy* knock out (Troy^KI/KI^) mice by crossing two heterozygous *Troy-CreER^T2^* knock in (Troy^WT/KI^) mice and confirmed a reduction in *Troy* expression (Figure S4A-B). We labelled Troy^WT/KI^ and Troy^KI/KI^ progenitor cells and compared their basal clone sizes after three months of lineage tracing to clones derived from Sox2^WT/KI^ mice. Clones derived from either Troy^WT/KI^ or Troy^KI/KI^ progenitors comprised more basal cells compared to clones derived from Sox2^WT/KI^ labelled progenitor cells (Figure 4A-B). In addition, Troy^KI/KI^ clone sizes were significantly larger than clones derived from Troy^WT/KI^ progenitors (Figure 4B).

**Figure 4.**
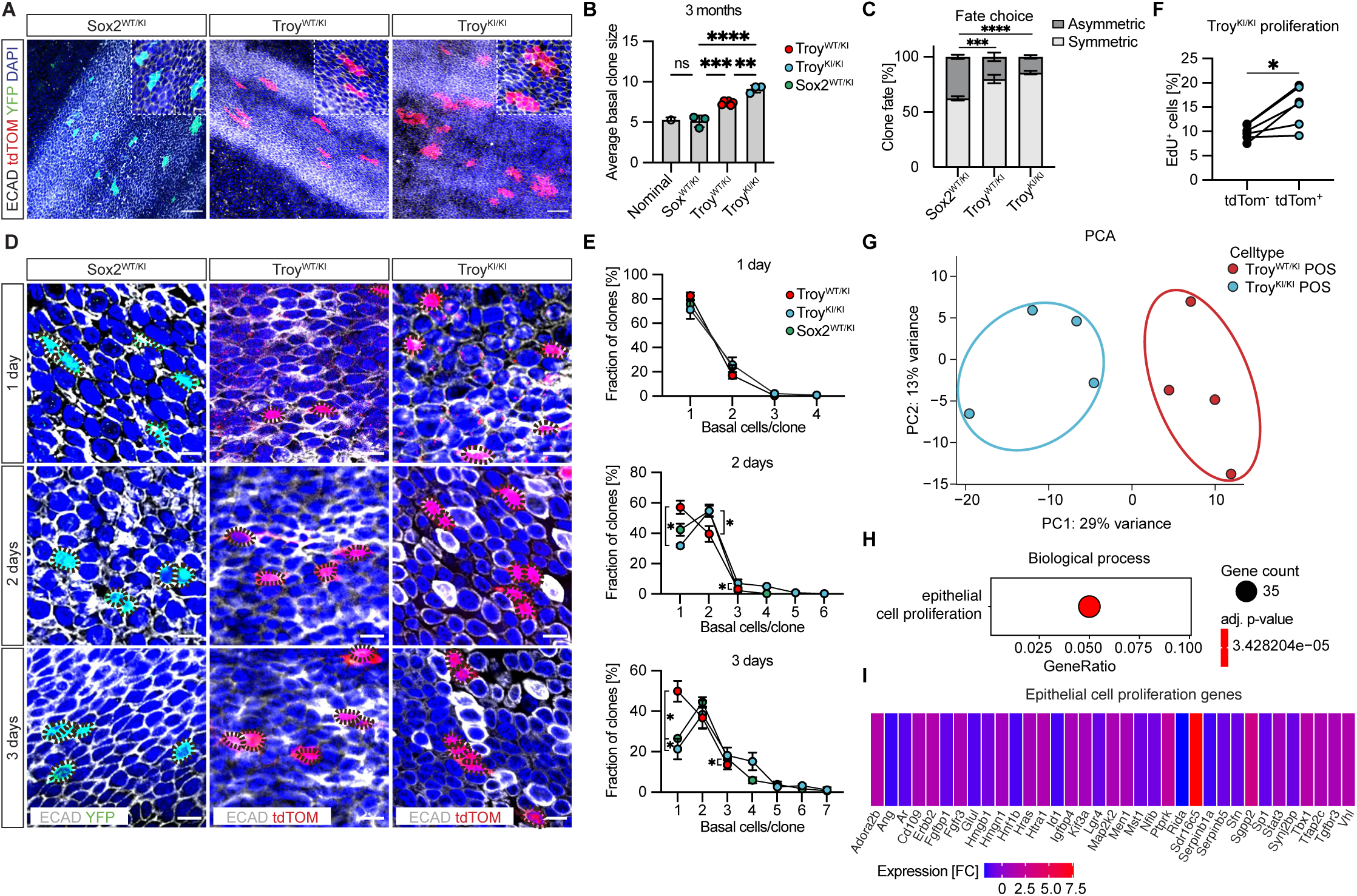
*Troy* negatively regulates progenitor cell proliferation. **(A)** Representative images of basal ECAD- and reporter-positive basal clone sizes after 3 months of lineage tracing in Sox2^WT/KI^, Troy^WT/KI^ and Troy^KI/KI^. **(B)** Average clone size at 3 months comparing Sox2^WT/KI^, Troy^WT/KI^ Troy^KI/KI^ experimental data with expected basal clone sized from the single-progenitor model (*r*=0.1, λ=2.9, γ=5.5) ^6^. **(C)** Quantification of symmetrically fated (2 basal or 2 suprabasal cell) or asymmetrically (1b1sb cell) clones in each mouse line at 2 days of tracing. **(D)** Representative images of clone size expansion during 1, 2 and 3 days of lineage tracing using Sox2^WT/KI^, Troy^WT/KI^ and Troy^KI/KI^ lines. **(E)** Basal clones size distribution at 1, 2 and 3 days of lineage tracing using Sox2^WT/KI^, Troy^WT/KI^ and Troy^KI/KI^ lines. **(F)** *In vivo* EdU incorporation is increased in tdTOMATO reporter-positive compared to reporter-negative basal cells in Troy^KI/KI^ mice. **(G)** Principal component analysis comparing the transcriptomes of tdTOMATO-positive progenitor cells FACS isolated from Troy^WT/KI^ and Troy^KI/KI^. **(H)** Gene ontology analysis reveals the differentially expressed genes to be associated with epithelial cell proliferation. **(I)** Relative expression of the 35 genes linked to epithelial cell proliferation in **(H)**. Data are represented as mean ± SD. ^∗^p< 0.05, ^∗∗^p<0.01, ^∗∗∗^p < 0.001. Statistical tests are: Ordinary one-way ANOVA (**B**, **C**), Wilcoxon matched-pairs signed ranked test (**F)** and multiple unpaired t-tests (**E)** (adjusted p-value < 0.05,). Scale bars are: (**D)** 10 μm; (**A)** 50 μm and inlets 20 μm. n=3-5 in all experiments.

To understand if fate choice was altered in Troy^KI/KI^ progenitors, we labelled and traced Sox2^WT/KI^, Troy^WT/KI^ and Troy^KI/KI^ progenitors for 1, 2 and 3 days. This short time frame enabled us to quantify all daughter cells generated upon division of labelled progenitor cells, thereby providing enough resolution to confirm fate choices potentially masked by altered proliferation kinetics in long-term tracing experiments. Comparing 1 and 2 days of lineage tracing (to capture clones after a single cell division) using CD49f and 3D imaging to discriminate between basal and suprabasal cells (Figure S4C-D), we focused on clones composed of two cells. All three transgenic lines demonstrated an increase in clones composed of one basal and one suprabasal (1b+1sb) cell comparing 1 and 2 days, although the overall frequency of clones with asymmetric fate (1b+1sb) was significantly higher in the Sox2^WT/KI^ compared to Troy^WT/KI^ and Troy^KI/KI^ progenitor cell derived clones (S4E-G). Comparing clone frequencies with symmetric (2b or 2sb) and asymmetric (1b1sb) fate 2 days after recombination confirmed that Sox2^WT/KI^ progenitor cells are more likely to generate asymmetrically fated daughter cells when compared to both Troy^WT/KI^ and Troy^KI/KI^ progenitor cells (Figure 4C). These observations indicate that both Troy^WT/KI^ and Troy^KI/KI^ progenitors favor symmetric fate choices when compared to Sox2^WT/KI^ progenitor cells.

### TROY negatively regulates progenitor cell proliferation

To detail if proliferation kinetics and fate choices of *Troy-*expressing progenitor cells are intertwined, we quantified basal clone sizes at 1, 2 and 3 days after tamoxifen. We found that clone sizes were strikingly similar at day 1, only to expand with genotype-specific dynamics at day 2 and 3 (Figure 4D-E). Whereas most clones derived from Troy^WT/KI^ either remained as single basal cells or formed small basal clones (2-3 cells), basal clones from both Sox2^WT/KI^ and Troy^KI/KI^ progenitors were larger (<7 cells) and few single basal cell clones remained after 3 days of tracing.

Since the average basal clone size in clones derived from Troy^KI/KI^ progenitors was increased when compared to both Troy^WT/KI^ or Sox^WT/KI^ after 3 months of tracing (Figure 4B), we injected Troy^KI/KI^ mice with EdU for 2 hours to directly probe how the absence of TROY affects progenitor cell proliferation. We found increased proliferation in Troy^KI/KI^ reporter-positive compared to neighboring *Troy* reporter-negative progenitor cells from the same animal or when comparing to the proliferation ratio of Troy^WT/KI^ progenitor cells (Figure 4F and S4H), demonstrating that TROY negatively affects esophageal progenitor cell proliferation.

### TROY impacts the transcriptional landscape of progenitor cells

To gain mechanistic insights into how TROY restricts progenitor proliferation, we isolated tdTOMATO-positive Troy^WT/KI^ and Troy^KI/KI^ progenitor cells 24 hours after tamoxifen injection and performed bulk RNA-sequencing (n=4 per genotype). Principal component analysis suggest that the two progenitor populations are distinct (Figure 4G) and differential gene expression analysis identified 761 genes to be enriched or downregulated in Troy^KI/KI^ when compared to Troy^WT/KI^ derived tdTOMATO-positive progenitor cells (Supplemental Table 5). Exploring Gene Ontology to understand the function of the differentially expressed genes identified epithelial cell proliferation as the only statistically significant altered biological process in the absence of TROY (Figure 4H-I)(Supplemental Table 6). Focusing on the 35 genes linked to epithelial proliferation, we identified differential expression of transcription factors as well as signaling receptors and effectors indicating that loss of TROY impacts both transcriptional regulation as well as the signaling landscape in esophageal progenitor cells.

### Commitment to differentiation is impaired in the absence of *Troy*

To understand if expression of *Troy* not only limits progenitor proliferation but also impinges on differentiation, we quantified the total number of basal and suprabasal cells labelled after 1, 2 and 3 days of lineage tracing and found that the basal-to-suprabasal ratio was significantly altered in Troy^KI/KI^ when compared to Troy^WT/KI^ mice (Figure 5A). Whereas clones derived from Troy^KI/KI^ were enriched for basal progenitor cells, Troy^WT/KI^ clones contained more suprabasal daughter cells. To assess if delamination, a proxy for differentiation, is affected by *Troy* we quantified the relative change in single labelled suprabasal cells in Troy^WT/KI^ and Troy^KI/KI^ esophagi comparing either 1-to-2 or 1-to-3 days of tracing and found a sustained increased number of suprabasal cells in Troy^WT/KI^ compared to Troy^KI/KI^ suggesting that Troy^KI/KI^ daughter cells are more often retained in the basal layer (Figure S5A).

**Figure 5.**
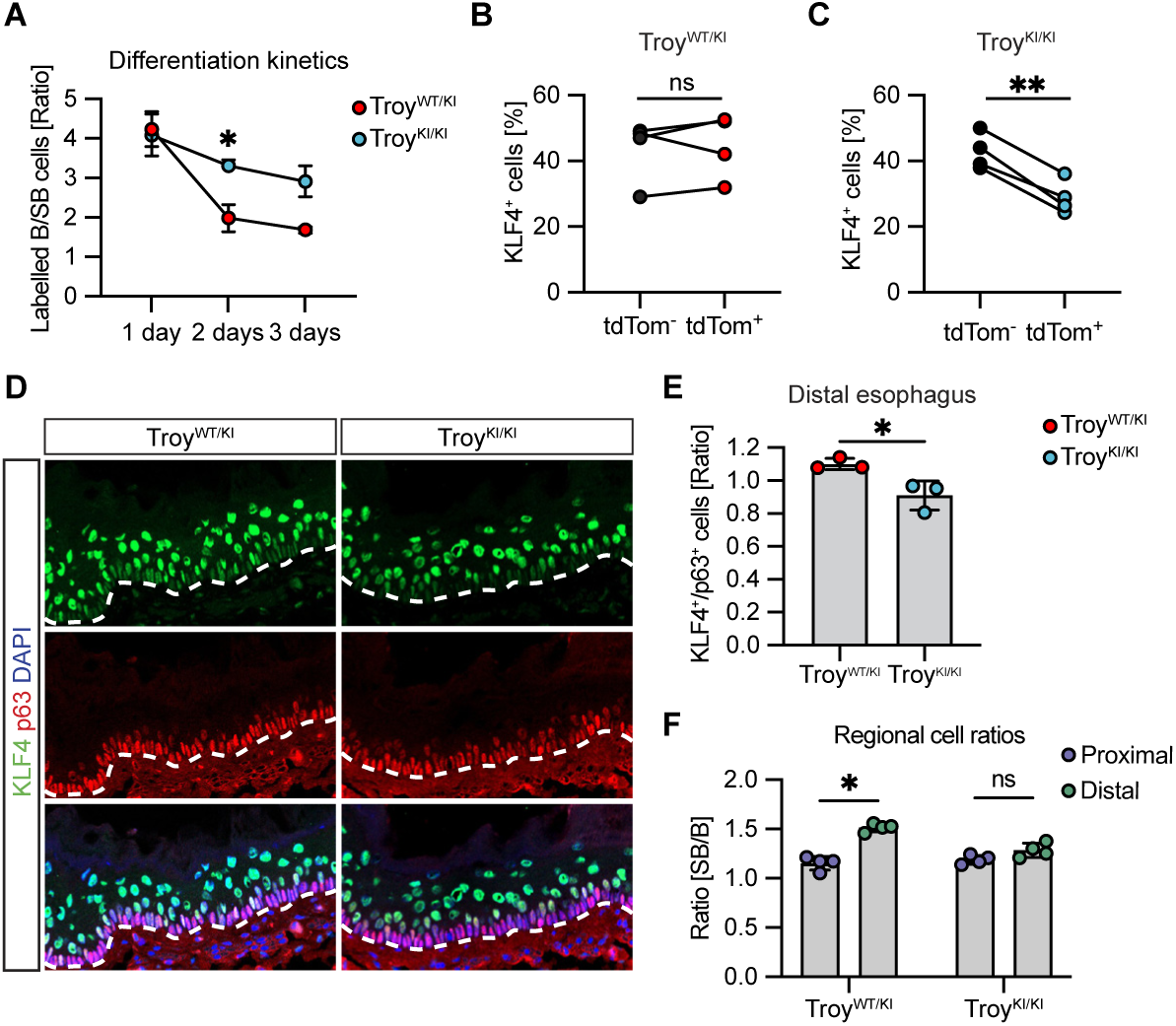
Commitment to differentiation is impaired in the absence of *Troy*. **(A)** Basal-to-suprabasal ratio of traced clones at 2 and 3 days is distinct comparing Troy^WT/KI^ and Troy^KI/KI^. **(B-C)** Quantification of immunoreactivity for KLF4 in tdTOMATO-negative and positive cells within the same Troy^WT/KI^ **(B)** and Troy^KI/KI^**(C)** esophagi. **(D-E)** Representative images and quantification of ratio of KLF4 to p63 positive cells in Troy^WT/KI^ and Troy^KI/KI^ distal regions enriched for *Troy*-positive progenitor cells. White dashed lines demark the epithelial stromal border. **(F)** Regional assessment of basal-to-suprabasal cell ratios in homeostasis. Data are represented as mean ± SD. ^∗^p< 0.05, ^∗∗^p<0.01, ^∗∗∗^p < 0.001. Statistical tests are: Two-sided unpaired t-test (**A**), Wilcoxon matched-pairs signed ranked test (**B** and **C),** Two-sided unpaired t-test **(E)** and multiple unpaired t-tests (**F)** (adjusted p-value < 0.05). Scale bars are: (**D)** 20μm. n=3-5 in all experiments.

KLF4 marks p63-positive basal cells committed to differentiation, as well as p63-negative differentiated suprabasal cells ^14^. We compared the KLF4 immunoreactivity in tdTOMATO-negative and positive cells in Troy^WT/KI^ and Troy^KI/KI^ mice 24 hours after tamoxifen. Whereas KLF4 expression was equally distributed in both tdTOMATO-positive and negative cells from the same animal in Troy^WT/KI^ esophagus (Figure 5B), KLF4 immunoreactivity was significantly reduced in tdTOMATO reporter-positive compared to reporter-negative cells in Troy^KI/KI^ esophagi (Figure 5C). To understand if the difference in KLF4 reactivity in tdTomato-positive clones correlated to differences in overall tissue differentiation dynamics, we analyzed the total number of p63 (basal) and KLF4-positive (suprabasal) cells in the distal epithelium, a region enriched for *Troy-*positive progenitor cells. We found a significant decrease in the KLF4/p63 ratio when comparing Troy^KI/KI^ to Troy^WT/KI^, indicating that the generation of suprabasal cells was impaired in the absence of TROY (Figure 5D-E).

Probing deeper into the transcriptional signature of Troy^KI/KI^ progenitor cells, we correlated the reduced number of KLF4-positive cells (Figure 5C) to *Klf4* expression and characterized additional esophageal differentiation markers (*Klf4*, *Krtdap*, *Krt13*, *Krt4* and *Id1*) which were all reduced in Troy^KI/KI^ compared to Troy^WT/KI^ progenitor cells (Figure S5B-C). In addition, we found that Troy^KI/KI^ progenitor cells express increased levels of *Itgb1*, *Itgb4, Itga3, Itga6* and *Itga7*, suggesting that availability of anchoring proteins could contribute to the basal retention of Troy^KI/KI^ progeny (Figure S5D). Collectively, these data suggest that in the absence of TROY, differentially fated progenitor cells fail to efficiently upregulate markers associated with commitment to differentiation and instead remain anchored to the basal membrane.

### Axial *Troy*-dependent regulation of esophageal tissue homeostasis

The proximal-to-distal gradient of *Troy*-expressing progenitor cells (Figure 1) suggest that aspects of esophageal homeostasis could be differentially regulated at an axial tissue level, functionally dependent on the frequency of *Troy-*positive progenitor cells. We quantified the total suprabasal-to-basal cell ratio in cross sections from proximal and distal esophagus and found that whereas the relationship between basal and suprabasal cells was 1.1-1.2 in proximal regions, the distal esophagus displayed significantly increased ratio of >1.5 suprabasal cells/basal cell in Troy^WT/KI^ mice (Figure 5F). Interestingly, Troy^KI/KI^ esophagi lost the regional basal-to-suprabasal cell profile and the distal Troy^KI/KI^ region (normally harboring most *Troy*-positive progenitors) was shifted towards a proximal cell ratio (Figure 5F), likely due to a distal expansion of the basal, and reduction of the suprabasal, cell population (Figure S5E-H). Collectively, these data not only confirm that TROY affects progenitor proliferation and differentiation but indicates that *Troy* expression is required to maintain regional esophageal tissue architecture.

### Troy^KI/KI^ proliferation dynamics are cell autonomous

*Troy* is expressed by a subset of fibroblasts in the stroma (Figure S1B). To understand if the basal expansion of Troy^KI/KI^ progenitor cells is caused by ablating TROY in the progenitor cells themselves, or because of secondary signaling effects due to lack of TROY in the stroma, we established organoids from Troy^WT/KI^ and Troy^KI/KI^ mice and assessed their size and marker expression. Both Troy^WT/KI^ and Troy^KI/KI^ organoids sustained long-term passaging (<24 passages) but whereas Troy^WT/KI^ organoids retained a normal round morphology, Troy^KI/KI^ organoids were larger and developed an irregular basal layer (Figure 6A and S6A). Troy^KI/KI^ organoids displayed a largely intact basal KRT5 expression, whereas immunoreactivity for the differentiation marker KRT13 was reduced (Figure 6A). Quantifying the overall cell number of individual organoids (Figure 6B) confirmed that Troy^KI/KI^ organoids were composed of more cells. Furthermore, more cells expressed p63 and KI67 in Troy^KI/KI^ compared to Troy^WT/KI^ organoids (Figure 6C-D) indicating that the basal proliferative progenitor population was expanded. We also recorded an increase in the number of KI67/KLF4 double-positive cells in Troy^KI/KI^ organoids, suggesting that commitment to differentiation was impaired (Figure 6E). While our data strongly suggest that TROY expression in esophageal progenitor cells act cell autonomously to restrict proliferation and enable differentiation, the function of TROY in neighboring fibroblasts is still unclear ^12^.

**Figure 6.**
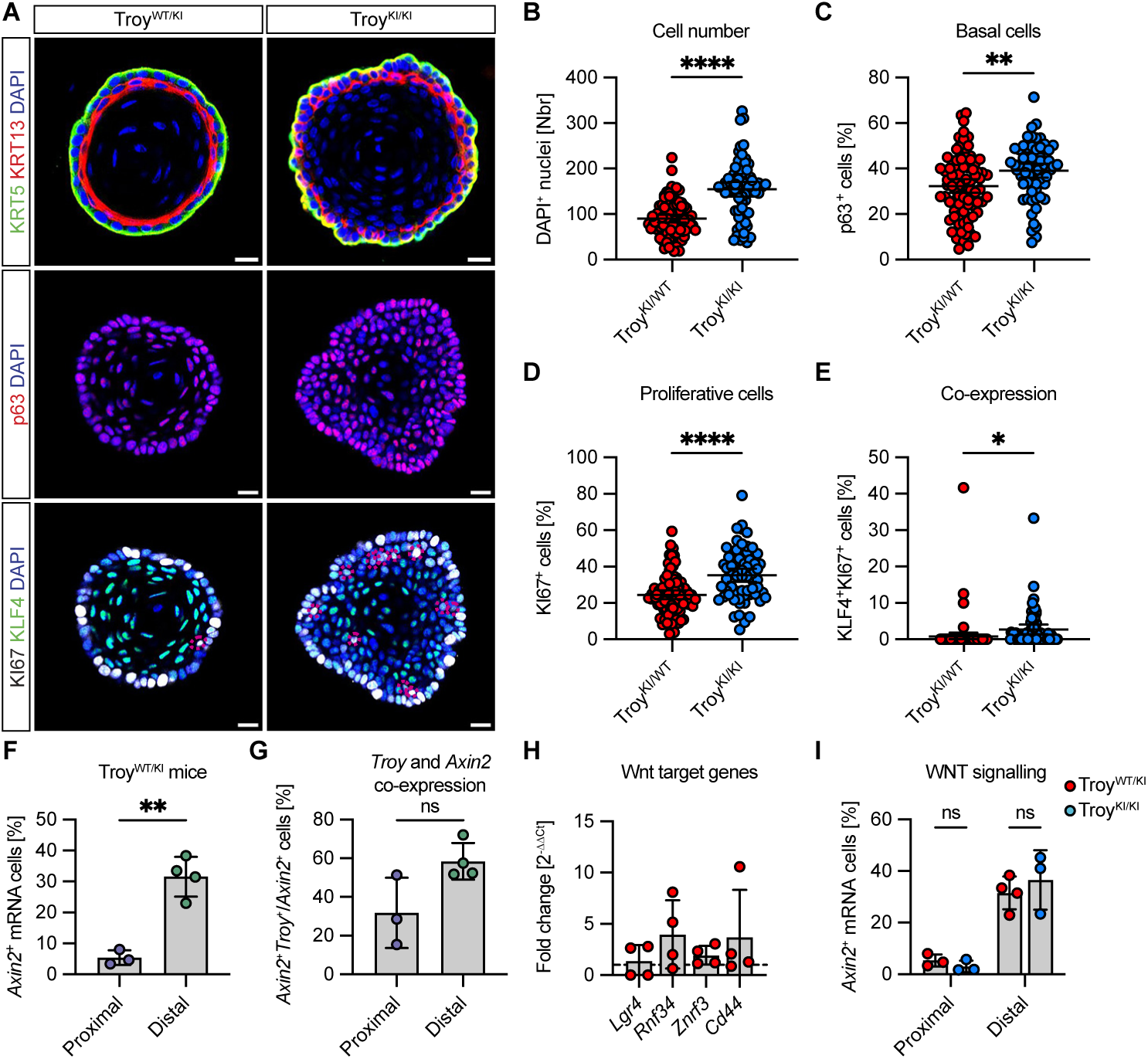
Troy^KI/KI^ proliferation dynamics are cell autonomous. **(A)** Representative images of Troy^WT/KI^ and Troy^KI/KI^ organoids after long-term passaging. Red dashed lines indicate cells co-expressing KLF4 and KI67. **(B)** Quantification of the total number of DAPI-positive nuclei in individual organoids. **(C)** Percentage of p63-expressing progenitor cells are increased in Troy^KI/KI^ when compared to Troy^WT/KI^. **(D)** Quantification of the percentage of KI67-positive, cycling, progenitor cells in organoids. **(E)** Troy^KI/KI^ organoids have an increased proportion of cells that co-express progenitor (KI67) and differentiation (KLF4) markers. **(F)** Quantification of *Axin2*-expressing basal cells comparing proximal and distal esophagus. **(G)** Percentage of *Axin2*-positive basal cells co-expressing *Troy*. **(H)** qPCR comparing WNT target genes in tdTOMATO-positive and tdTOMATO-negative *in vivo* isolated progenitor cells. **(I)** Comparison of *Axin2* mRNA expression in proximal and distal esophagus from Troy^WT/KI^ and Troy^KI/KI^ mice. Data are represented as mean ± SD. ^∗^p< 0.05, ^∗∗^p<0.01, ^∗∗∗^p < 0.001. Statistical tests are: Two-sided unpaired t-test (B-E), Two-sided paired t-test (F and G), multiple unpaired t-tests (adjusted p-value < 0.05) (I). Scale bars are: (**A)** 20μm.

WNT signaling affects progenitor proliferation and renewal ^34^, and TROY is both a reported WNT target gene ^35^ and negative regulator of WNT signaling ^16^. To characterize active WNT signaling, we assessed the expression of *Axin2* in the basal layer and found that the *Axin2* expression was increased in the distal compared to proximal esophagus (Figure 6F and S6B). Quantifying the number of progenitors co-expressing *Troy* and *Axin2* revealed that up to 60% of *Axin2*-positive progenitor cells also expressed *Troy* (Figure 6G). Using qPCR to compare the expression of additional WNT target genes *Lgr4*, *Rnf34*, *Znrf3* and *Cd44* ^36-38^ in FACS isolated *Troy*-positive and *Troy*-negative progenitor cells revealed that WNT signaling activity correlates to *Troy* expression (Figure 6H). To understand if WNT signaling activity is altered in the absence of TROY ^16^, we assessed *Axin2* expression in Troy^WT/KI^ and Troy^KI/KI^ and found that the number of *Axin2*-positive progenitor cells was not significantly different when comparing Troy^WT/KI^ and Troy^KI/KI^ (Figure 6I and S6B). These data indicate that while *Troy* is a possible WNT target gene in the esophagus, TROY does not act to inhibit canonical WNT signaling.

### *Troy*-positive progenitor cells are activated in response to atRA

Upon tissue challenge, it is pertinent to quickly generate new basal cells to restore tissue integrity and homeostasis. Our transcriptional profiling identifies the immediate early family of transcription factors to be enriched in *Troy*-expressing progenitor cells (Figure 2), indicating that *Troy* expression could mark a cell state primed to respond to environmental changes.

All-trans retinoic acid (atRA) acts to enhance differentiation and delamination of basal cells ^2,7,39^, subsequently inducing progenitor proliferation in order to regenerate the esophageal epithelium. We injected atRA every second day over the course of nine days and administered tamoxifen once, 24 hours before sacrifice (Figure 7A-B). Quantification revealed that the number of Troy^WT/KI^ tdTOMATO-expressing basal cells was diminished when compared to control esophagi (Figure 7C-D). Additionally, we found that the relative proliferation of remaining Troy^WT/KI^ reporter-positive and -negative progenitor cells were altered upon atRA administration. Normally less proliferative Troy^WT/KI^ reporter-positive progenitors (Figure 7E) incorporated significantly more EdU than *Troy-*negative progenitor cells after atRA treatment (Figure 7F), demonstrating that *Troy*-positive progenitor cells are preferentially activated when homeostasis of the esophageal epithelium is challenged (Figure 7G).

**Figure 7.**
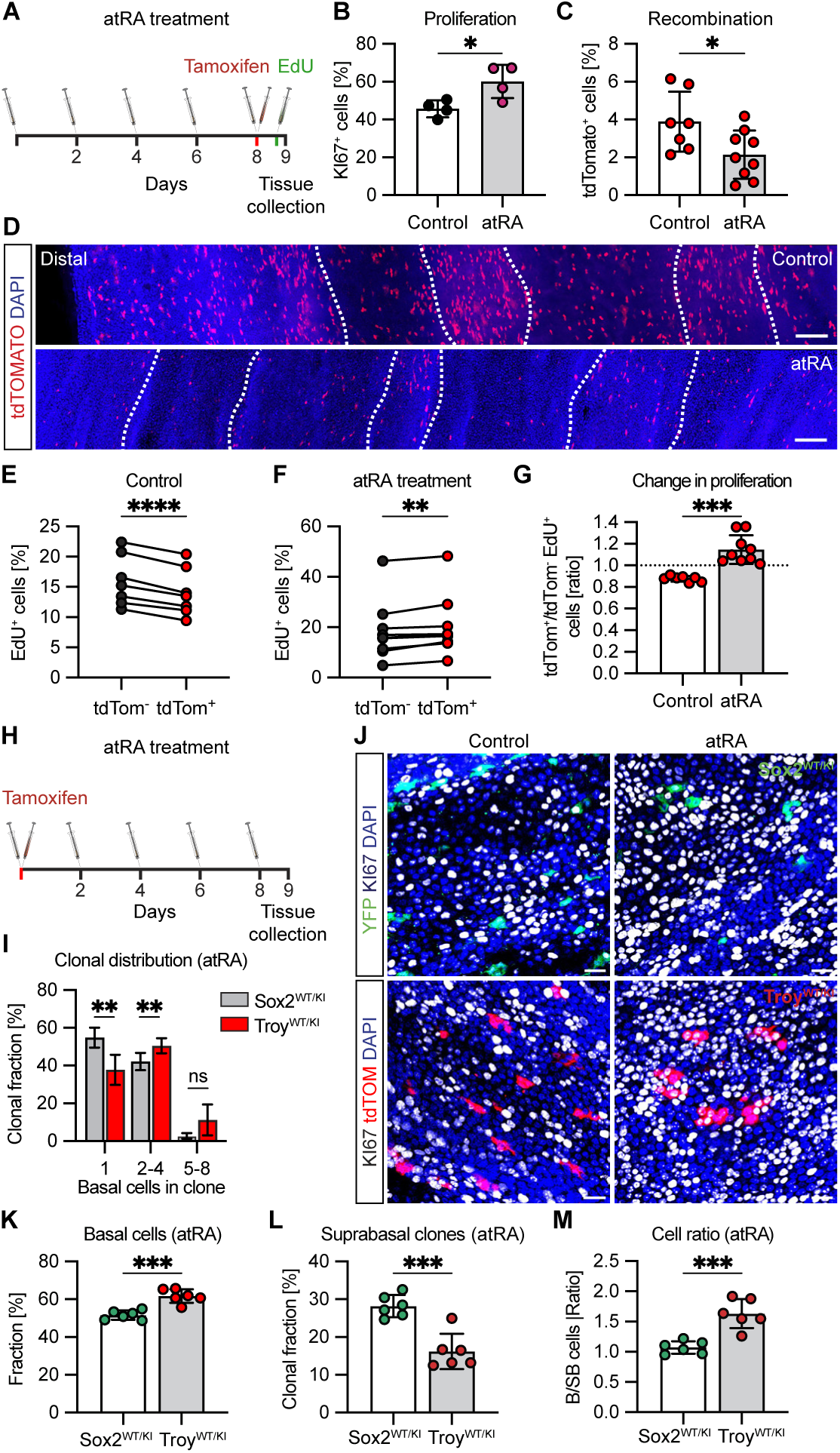
*Troy* is dynamically regulated upon tissue stress. **(A)** Schematic representation of atRA administration. Tamoxifen is administered at day 8 and EdU 2 hours before sacrifice. **(B)** All-trans retinoic acid (atRA) induces epithelial proliferation as judged by KI67 immunoreactivity. **(C-D)** The percentage of tdTOMATO reporter-positive progenitor cells is reduced in response to atRA. **(E-F)** EdU incorporation in control and atRA treated esophagi. **(G)** Relative proliferation comparing tdTOMOTO-negative and tdTOMATO-positive cells normally and upon atRA treatment **(H)**. Tamoxifen is administered at day 1 to enable labelling of *Troy*-positive progenitor cells before atRA administration. **(I)** Basal clone size distribution comparing Sox2^WT/KI^ and Troy^WT/KI^ after treatment with atRA. **(J)** Immunofluorescence demonstrating increased KI67 in atRA treated compared to control esophagi, in both Sox2^WT/KI^ and Troy^WT/KI^. Reporter expression (YFP or tdTOMATO) reveal clone size distribution in both lines comparing control and atRA treatment. **(K-L)** Fraction of all labelled cells which are basal **(K)** or percentage of suprabasal, floating, clones (without basal cell contribution) **(L)**, comparing Sox2^WT/KI^ to Troy^WT/KI^ after atRA treatment. **(M)** Basal-to-suprabasal ratio comparing the total number of all basal and suprabasal cells in Sox2^WT/KI^ and Troy^WT/KI^ clones after atRA treatment. Data are represented as mean ± SD. ^∗^p< 0.05, ^∗∗^p<0.01, ^∗∗∗^p < 0.001, ^∗∗∗∗^ p < 0.0001. Statistical tests are: Two-sided unpaired t-test (**A**, **F**, **J**, **K** and **L**), Wilcoxon matched-pairs signed ranked test (**D** and **E**). Scale bars are: 200μm (**C**) and 20μm (**I**). n=4-9 in all experiments.

### *Troy*-derived clones are refractory to atRA induced differentiation

To address if *Troy*-positive progenitors are lost when the esophageal epithelium is challenged (Figure 7C-D), we induced reporter expression in both Troy^WT/KI^ and Sox2^WT/KI^ mice at the first day of atRA treatment (Figure 7H). Visualization and quantification of KI67 confirmed that atRA increased the overall proliferation rates in both lines (Figure 7J and S7A-B). The frequency of Sox2^WT/KI^ or Troy^WT/KI^ derived basal clones did however not change when comparing control to atRA treated esophagi within lines (Figure S7C-D), indicating that the reduction of Troy^WT/KI^ reporter-positive progenitors seen at day 9 of atRA treatment is likely caused by silencing of *Troy* expression rather than selective loss of *Troy-*positive progenitor cells.

To understand if atRA-induced activation of *Troy-*positive progenitor cells resulted in increased basal contribution, we analyzed the basal clone size distribution comparing Sox2^WT/KI^ and Troy^WT/KI^ atRA treated and control esophagi. In general, the overall basal clone sizes were not significantly different between control and atRA within the same line, although larger clones (5-8 basal cells) were found in atRA treated but not control Troy^WT/KI^ esophagi (Figure S7E-F), supporting the idea that *Troy-*positive progenitor cells are preferentially activated during esophageal tissue stress. Comparing Sox2^WT/KI^ and Troy^WT/KI^ basal clone sizes after atRA revealed that Troy^WT/KI^ clones were larger (reduction in 1 basal cell clones coupled with an increase in 2-4 and 5-8 basal cell clones) than clones derived from Sox2^WT/KI^ progenitor cells (Figure 7I-J). These data suggest that *Troy-*positive progenitors produce larger basal cell clones compared to *Troy-*negative progenitor cells during esophageal tissue stress.

Quantification of the total number of labelled basal and suprabasal cells showed that the Sox2^WT/KI^-derived clone composition shifted towards containing more suprabasal cells (Figure S7G-I), whereas the cell composition in Troy^WT/KI^-derived clones was unaltered when comparing control and atRA treated mice (Figure S7J-L). Comparing the cell composition of Sox2^WT/KI^ to Troy^WT/KI^ clones after atRA confirmed that the basal cell fraction was significantly increased whereas the number of suprabasal (floating) clones were reduced in Troy^WT/KI^ when compared to Sox2^WT/KI^ (Figure 7K-L). The overall basal to suprabasal cell ratio was also higher in Troy^WT/KI^ compared to Sox2^WT/KI^, indicating that Troy^WT/KI^ cells were less prone to differentiate upon atRA treatment (Figure 7M). These data suggest that in the presence of atRA, *Troy-*negative (Sox2^WT/KI^) progenitors generate clones prone to delamination whereas Troy^WT/KI^ daughters are retained in the basal layer, thereby confirming the co-existence of functionally non-equivalent progenitor populations in the esophageal epithelium.

## Discussion

Progenitor cells maintain tissue homeostasis and are able to adapt their behavior in response to tissue injury ^40^. Here we characterize a subset of esophageal progenitor cells marked by *Troy* expression and delineate the contribution of *Troy-*positive progenitor cells to the esophageal epithelium in homeostasis and tissue stress. Although the esophagus is traditionally seen as a uniform tube, we identify a proximal-to-distal gradient of *Troy-*positive progenitor cells. In addition, we find prominent enrichment of *Troy* expression around the tube circumference in the distal esophagus (Figure 1) disclosing local epithelial heterogeneity likely to extend beyond epithelial *Troy* expression.

Detailing progenitor characteristics reveals that although *Troy-*expressing progenitors are less proliferative than neighboring *Troy-*negative progenitors, they are more than twice as good at forming organoids (Figure 2), suggesting that *Troy-*positive progenitor cells are able to undergo renewing divisions. Although using an unbiased (Sox2^WT/KI^) labelling strategy faithfully recapitulates previous work^7^ (Figure 3), tracing of *Troy*-positive progenitor cells deviates from the suggested progenitor cell behavior. Juxtaposing the *Troy*-derived data with quantitative lineage tracing derived from unbiased Sox2^WT/KI^ basal cell labelling suggest that *Troy* progenitor cells are more likely to generate daughter cells of the same fate (differentiated-differentiated or progenitor-progenitor), compared to the low symmetric fate probabilities of *Troy*-negative progenitors (Figure 3). Despite changes in fate choice, our data show that *Troy*-positive progenitors compete neutrally with neighboring *Troy*-negative progenitor cells. Furthermore, the balanced expansion and loss of clones suggest that *Troy*-derived clones fulfil all characteristics of neutral drift, where the likelihoods of generating either of the two symmetric fate outcomes (two progenitor or two differentiated daughter cells) are equally balanced within the *Troy*-positive as well as the *Troy*-negative progenitor population. Our data suggests that two neutrally competing progenitor populations with distinct fate probabilities act in concert to mediate esophageal tissue homeostasis.

Whereas fate probabilities are distinct in *Troy*-positive and *Troy*-negative progenitor cells during esophageal homeostasis (Figure 3) our work indicate that TROY is itself dispensable for this fate choice. The molecular cues impacting and diversifying fate choices in the esophageal epithelium are yet unknown. We detect the presence of *Axin2*-positive progenitor cells in the esophagus indicating that canonical WNT signaling is active in the epithelium and could act on fate choice. However, the low *Axin2* mRNA-levels and the distal circumferential pattern of alternating regions of high and low abundance of *Troy*-positive progenitor cells suggest additional mechanisms in play, including local stromal gradients of signaling molecules or mechanical force dynamics induced upon expansion of the esophageal tube when feeding.

Esophageal tissue regeneration is associated with changes in progenitor proliferation rates and commitment to differentiation - but not fate choice ^7^. We show that the esophageal epithelium silence *Troy* expression in response to atRA stress (Figure 7) enabling symmetrically fated progenitor cells to robustly proliferate and contribute daughter cells to the basal layer regeneration. Strikingly, the stress-induced changes in *Troy*-positive progenitor cell behavior are mirrored by genetically ablating TROY. Troy^KI/KI^ progenitor cells are highly proliferative and generate daughter cells refractory to differentiation even in the absence of tissue challenge, demonstrating how controlling TROY expression could function as a molecular rheostat restricting proliferation normally when TROY expression is high, only to facilitate a rapid and coordinated basal layer expansion when *Troy* is silenced in response to tissue challenge (Figure 7).

We report the co-existence of two functionally non-equivalent, but neutrally competitive, progenitor populations in the esophageal epithelium.

## Limitations of the study

We perform and compare lineage tracing results from multiple transgenic mouse lines to evaluate the behavior of two cell populations in the esophagus. It is difficult to completely rule out how the results are impacted by different genetic backgrounds. Dual labelling of two or more esophageal cell populations in the same mouse line remains to be reported.

## Acknowledgements

This study was supported by ERC Starting Grant (TroyCAN 851241), Cancerfonden (19 0007-Pj), SFO StratRegen 2023/24 Junior Grant and Vetenskapsrådet (2023-02743). MG is a Cancerfonden Senior Investigator. We are grateful for technical assistance from Biomedicum Flow Cytometry Core Facility for FACS isolation and the Biomedicum Imaging Core facility (BIC) for confocal microscopy. We acknowledge support from the National Genomics Infrastructure in Stockholm funded by Science for Life Laboratory, the Knut and Alice Wallenberg Foundation and the Swedish Research Council, and SNIC/Uppsala Multidisciplinary Center for Advanced Computational Science for assistance with massively parallel sequencing and access to the UPPMAX computational infrastructure. We thank Kylie Hin-Man Mak for technical assistance with organoid cultures.

## Author contributions

DG and EE performed the experiments. DG, MS, MW, QD and PG analyzed the data. DG, PG and MG designed the experiments and drafted the manuscript. All authors have proof-read and approved the final version of the manuscript.

## STAR Methods

### RESOURCE AVAILABILITY

#### Lead contact

Further information and reasonable requests for resources and reagents should be directed to and fulfilled by the lead contact, Maria Genander

#### Materials availability

This study did not generate new unique reagents.

#### Data and code availability

Single-cell as well as bulk RNA-sequencing raw data are publicly available (GEO accession numbers GSE255402 and GSE255401). Original code used for modelling of fate choice can be found here (https://doi.org/10.5281/zenodo.10781132). Any additional information required to reanalyze the data reported in this paper is available from the lead contact upon reasonable request.

### EXPERIMENTAL MODEL AND STUDY PARTICIPANT DETAILS

All animals were free from mouse viral pathogens, ectoparasites and endoparasites and mouse bacteria pathogens. Mice were kept with the following light/dark cycle: dawn 6:00-7:00, daylight 7:00-18:00, dusk 18:00-19:00, night 19:00-6:00 and housing to a maximum number of 5 per cage in individually ventilated cages (IVC sealsafe GM500, tecniplast). General housing parameters such as relative humidity, temperature, and ventilation follow the European convention for the protection of vertebrate animals used for experimental and other scientific purposes treaty ETS 123, Strasbourg 18.03.1996/01.01.1991. Monitoring of husbandry parameters is done using ScanClime® (Scanbur) units. Cages contained hardwood bedding (TAPVEI, Estonia), nesting material, shredded paper, gnawing sticks and card box shelter (Scanbur). The mice received a regular chow diet (either R70 diet or R34, Lantmännen Lantbruk, Sweden). Water was provided by using a water bottle, which was changed weekly. Cages were changed every other week. All cage changes were done in a laminar air-flow cabinet. All animal experiments were approved by Stockholms djurförsöksetiska nämnd (ethical permit no N243/14, N116/16 and 14051-2019). Troy-CreER^T2^ and Confetti mice were previously described ^18,41^ and generously provided by Hans Clevers, Hubrecht Institute. Sox2-CreER^T2^ and PdgfRa-H2BeGFP lines are available at JAX (Strain #017593 and #007669 respectively) ^42,43^. Experiments were started when mice were between 10 and 14 weeks of age.

### METHOD DETAILS

#### Animal Experiments

Mouse littermates of the same sex were randomly assigned to experimental groups. All experiments were performed on male and female mice. Except ear clipping for genotyping purposes no previous procedures were performed on experimental animals. Studies were not blinded. Three or more mice per experimental group was used to ensure statistical power. No data points or animals were excluded from the final data representation.

##### Genotyping

Ear or tail biopsies were lysed overnight (o/n) at 55°C in DirectPCR lysis reagent (BioSite) in the presence of 0.1mg/mL Proteinase K (Thermo Scientific). Lysis was stopped by heat inactivation (45min, 85°C). Taq Polymerase (5 U/µL, final concentration: 0.05U/µL) was used for amplification in PCR buffer supplemented with 0.08mM MgCl2, 0.2mM dNTPs (all Invitrogen) and 0.4µM primers. PCR products were analyzed with 2% agarose gel.

##### Tamoxifen and EdU labelling

For the recombination of Troy-CreER^T2^:R26tdTomato, 200ul of 1% Tamoxifen (Sigma, #T5648) in ethanol was administered once topically on the back skin 24 hours before sacrificing the mice. To recombine Sox2-CreER^T2^:R26YFP, 200ul of 0,01% Tamoxifen (Sigma) in ethanol was administered topically on the back skin. The Tamoxifen dose (0,01%) was titered to reflect the recombination efficiency of the Troy-CreER^T2^ line. Recombination in Sox2-CreER^T2^:R26YFP mice is uniformly distributed along the esophageal axis. Topical tamoxifen application has been extensively used before in epithelial research and is sufficient for efficient recombination ^44-46^. Due to the low recombination efficiency of the Confetti locus, 100ul of 5% Tamoxifen was injected intraperitoneally on 5 consecutive days to Troy-CreER^T2^ : R26Confetti mice. All lines were controlled for leakage of recombination by assessing whole mount staining from untreated CRE+ mice (see also Figure S2E for FACS quantification). To investigate circadian *Troy* expression, mice were topically treated with 200ul of 1% Tamoxifen (Sigma, #T5648) in ethanol 24 hours before intraperitoneal EdU injections, respectively (Day: 7AM; Night: 7PM). Mice were sacrificed 1 hour after EdU injections (Day: 8AM; Night: 8PM). For all other experiments, dividing cells were labelled by injecting 100ul of 1mg/ml of EdU intraperitoneally between 6 and 7AM. Mice were sacrificed 2 hours later, unless otherwise mentioned in the figure legend.

##### All-trans Retinoic Acid treatment

100ul (4mg/ml in DMSO) all-trans retinoic acid (R2625-100MG, SigmaAldrich) or DMSO was injected intraperitoneally every second day for 9 days. On the last injection day mice were additionally topically treated with 200ul Tamoxifen (1% W/V) (T5648-5G, SigmaAldrich) in ethanol. 2 hours prior to euthanizing mice were injected intraperitoneally with 100ul of 1mg/mL EdU (A10044, ThermoFisher) in PBS. Mice were sacrificed using CO2 and the esophagi dissected. Esophagi were prepared for whole mounts, or directly fixed in 4% formaldehyde (252549-1L; SigmaAldrich).

#### Whole mount preparations and labelling

To prepare whole mount staining of the entire esophagus, esophagi were dissected and the muscle layer surrounding the esophagus was mechanically removed using forceps. Subsequently, the esophagus was longitudinally cut open and cut into approximately 5 mm pieces. Pieces were incubated for 2-3 hours in 5 mM EDTA in PBS at 37°C and the epithelium peeled away from the underlying tissue ^47^. Afterwards, the epithelium was fixed in 4% formaldehyde (FA) in PBS for 30 min. For staining, wholemounts were blocked for around 1 hour in staining buffer (2,5% normal goat serum, 2,5% normal donkey serum, 1% BSA, 0,5% Triton X-100 in PBS). Primary antibodies were incubated in staining buffer overnight. Subsequently, pieces were washed for 2 hours with 0,2% Tween-20 in PBS exchanging the washing buffer every 30min. Secondary antibodies were incubated in staining buffer for 3 hours at room temperature in the dark followed by another 2 hours washing procedure. Next, pieces were incubated overnight in DAPI (250ng/mL) in PBS. Lastly, pieces were spread out on object slides and mounted. Images were obtained with either a Zeiss LSM900, Zeiss LSM980 confocal microscopes or a Zeiss AxioImagerZ2 microscope.

#### Antibody staining

Tissue samples used in this study were fixed in 4% Formaldehyde (Sigma) and embedded in OCT. Samples used for immunofluorescence staining were sectionned at a thickness of 10µm, blocked with donkey serum (2,5%), goat serum (2,5%), and BSA (1%) and permeabilized with 0,3% Triton X-100. When EdU was analyzed, the samples were treated with click chemistry prior to the antibody labelling (Click-iT, Thermo Fisher). Antibodies used for immunofluorescence staining are listed below. KRT5 (Biolegend, #905901, 1:1000), KRT4 (Abcam, #ab9004, 1:500), KRT13 (Abcam, #ab92551, 1:500), KRT14 (Acris Antibodies, #BP5009, 1:1000), KLF4 (Abcam, #ab214666, 1:400), SOX2 (Abcam, #ab92494, 1:500), KI67 (eBioscience, #14-5698, 1:1000), CD49f (eBioscience, #14-0495-82, 1:500), p63 (Abcam, #ab735, 1:100), CCNA2 (Abcam, #ab181591, 1:400), E-Cadherin (CST, #3195, 1:1000).

#### Isolation of esophageal progenitor cells

For isolation of esophageal progenitor cells, the muscle layer was removed from the submucosa and the remaining esophagus incubated in 0.5 mg/ml Thermolysin (Sigma) at 37°C for 15 min^47^. After incubation, the submucosa and the epithelium were separated. To obtain a single-cell suspension, the epithelium was minced and incubated in 0.25mg/ml DNaseI (10104159001, SigmaAldrich) and 0.25mg/ml LiberaseTM (5401127001, SigmaAldrich) at 37°C for 1 hour and resuspended every 15min. After incubation, the cells were filtered through a 40μm filter and cells were stained with CD324-PE/Cy7 (Biolegend, #147310) and CD104-A647 (Biolegend, #23608) for 30min on ice. Using FACS, the progenitor cells (CD324+CD104+) were separated and collected from the cell suspension. When progenitor cells were FACS isolated for organoid cultures, both reporter-negative and -positive cells were selected from the CD324+CD104+ basal progenitor population.

#### Esophageal organoid cultures

Esophagi were dissected and the outer muscle layer mechanically removed and esophageal epithelial cells were isolated as previously described ^47^. All epithelial cells were gated on CD324+CD104+ to enrich for basal cells. Subsequently, 2,500 basal cells were resuspended in Matrigel (356231, FisherScientific) medium mixture. Matrigel domes were plated on preheated cell culture plates (Sarstedt). Domes were allowed to dry for about 20min at 37°C. Organoid medium was added to cover the domes (DMEM/F-12 (11320033, Gibco), 1XPenStrep (15140122, Gibco), 1XB27 supplement (17504044, Gibco), 1XGlutaMAX^TM^ supplement (35050061, Gibco), 1mM N-Acetyl-cysteine (A9165-25G, SigmaAldrich). Medium was supplemented with 200ng/ml recombinant mouse R-spondin 1 (3474-RS-250, Biotechne), 100ng/ml murine Noggin (250-38, Peprotech), and 50ng/ml murine EGF (2028-EG-200, R&D systems). For the first two days of culture additionally 10µM Y-27632 dihydrochloride was added (Y0503-5MG, SigmaAldrich). Medium was replaced every second day.

#### Single-cell RNA-Sequencing

##### Sample preparation and library construction

Esophageal single cell suspension pooled from 2 male and 2 female wild type mice was obtained. To enrich for basal cells only CD104^+^CD49f^+^ cells were selected. The 10X Chromium platform was utilized for profiling of gene expression at the single-cell level. RNA isolation and library preparation were conducted by NGI-Sweden (National Genomic Infrastructure) at SciLifeLab, Stockholm. Samples were sequenced using a NovaSeq6000 platform (Illumina). The sample was run on two lanes of the same flowcell. 10X’s CellRanger v3.0.1 pipeline was used for pre-processing of the data by NGI-Sweden.

##### Single-cell RNA-seq data analysis

The reads were mapped to mm10 reference genome with Cell Ranger pipeline (version 3.0.1) with default parameters. We first conducted quality control. Only cells which met the following criteria were included for further analysis: more than 2,000 read counts in more than 1,000 genes and having less than 25% mitochondrial genes. 9,309 out of 11,164 cells were kept. Genes that were not expressed were filtered out. The UMI count matrix was analyzed by RaceID v0.1.3 ^28^ to identify cell clusters. In general, the major cluster number was determined by saturation point of the within-cluster dispersion and outliers in the initial k-medoids clusters. Firstly, we ran the algorithm using cln=4. Based on marker genes, the clusters assigned to endothelial cells, immune cells, myofibroblasts and blood cell were removed. Then, we ran a second round of clustering with remaining 8,767 cells using cln=3, outlg=20. The t-SNE representation was calculated by comptsne function using perplexity=200, and the Fruchterman-Rheingold graph layout was calculated by compfr function using knn=100. We used the function diffexpnb, which is similar to DESeq2 algorithm, to identify the differentially expressed genes in each cluster. GO enrichment was conducted on biological process subontologies. We used enrichGO function from ClusterProfiler package v3.11.1 ^48^, with simplify cutoff 0.7 and adjusted *p-value* 0.05. The annotation database is org.Mm.eg.db v3.7.0 (Carlsson (2019). *org.Mm.eg.db: Genome wide annotation for Mouse*. R package version 3.8.2). To understand if gender influenced cell clustering, we calculated the expression level of the Y chromosome for each cell. If the read count sum of Y chromosome was more than 1 then the cell was annotated as male, otherwise as female. To perform correlation heatmap, we picked the top 100 highly variable genes from each cluster and calculated the average expression value of each cluster. Then we performed Spearman correlation analysis to evaluate the similarity between clusters. The results were visualized by heatmap.

#### Bulk RNA-Sequencing

Single cell suspension from 4 separate mice (2 male and 2 female) was obtained according to the above-described procedure. Cells were isolated by FACS gating on CD324+,CD104+ and tdTOMATO+. RNA was isolated and libraries prepared according to Picelli *et al* ^49^. Samples were sequenced using a NovaSeq6000 platform (Illumina) on a single lane with 2x50bp reads. A quality check (Fast QC/0.11.5) was run on raw sequencing reads. Raw sequencing reads were processed to obtain counts per genes for each sample.

##### Bulk RNA-sequencing data analysis

The reads were aligned to the mm10 reference genome (GRCm38, vM25 annotation). After indexing and sorting, feature counts were obtained (samtools v1.10). Data was analyzed using DESeq2 (v 1.30.0) using tdTOMATO^+^ cells of Troy^WT/KI^ and Troy^KI/KI^ mice as a contrast. Only genes with a total count >10 were kept for subsequent analysis. Regularized logarithm transformed data was used for data visualization. clusterProfiler (v 3.18.0) was utilized for GO enrichment analysis with a simplified cutoff of 0.5.

#### Long term lineage tracing

All quantifications of clone sizes after lineage tracing were performed under live acquisition mode in esophageal whole mounts imaged using a confocal microscope. Only epithelium from the proximal and middle esophagus was used to avoid areas of high *Troy*-expression and reduce the risk of including fused clones. An average of 300 clones were quantified per animal and time point. 3-5 animals were used for each time point.

#### Mathematical modelling

All modelling is based on the published stochastic fate model ^6,7^ assuming that progenitor cell divisions over time generate equal number of new progenitor (P) and differentiated (B) daughter cells in order to maintain tissue homeostasis. The model is a continuous time Markov process model that captures time evolution of progenitor cells.

The model is

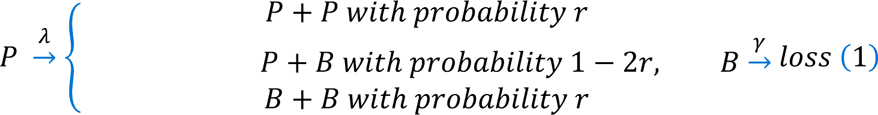

The division of the P cells is governed by an exponential (Poisson) time distribution with average rate *λ*. It will create either two P cells (with probability *r*) two B cells (with probability *r*) or one P and one B cell (with probability 1 − 2*r*). The stratification of B cells is governed by an independent exponential (Poisson) time distribution with average rate *γ*. When a B cell stratifies to the supra basal cell layer, we refer to it as a loss. We will here summarize how this model has been used in our analysis. The approach is very similar to the one in ^7^ and we refer the reader to the supplementary material of that paper for a more thorough discussion of the validity of the model.

##### Clone size distributions

To compute the clone size distributions at any given time *t*, we use the approach described in ^7^ with minor straightforward modifications arising from the fact that we consider only the basal layer cells in the clones. We summarize the approach here with the necessary modifications. Let *p*_*nm*_(*t*) denote the probability of having *n* progenitor cells (P) and *m* differentiated cells at time *t*. The probabilities generated from the underlying continuous time Markov process are governed by the Master equation (in which the time dependence of the probabilities *p*_*nm*_ has been suppressed)

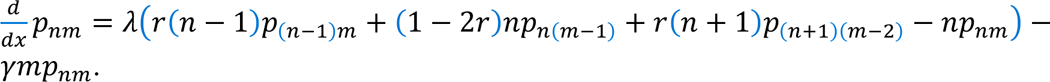

We initialize at time *t* = 0 with *p*_)-_(0) = 1 and *p*_*nm*_(0) = 0 for all other combinations of *n* and *m*, which corresponds to initializing with one progenitor cell (P) at time 0. This set of coupled ordinary differential equations (ODEs) is solved via the generating function, which is defined as *G*(*x*, *y*, *t*) = ∑_*nm*_ *p*_*nm*_(*t*)*x*^*n*^*y*^*m*^ for each *x*, *y*, *t* and satisfies the partial differential equation (PDE)

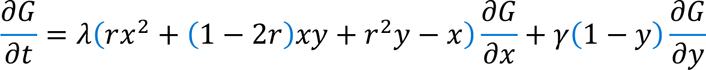

The initial condition is *G*(*x*, *y*, *t*) = *x*, corresponding to starting with one progenitor cell at time 0. This partial differential equation is solved via the method of characteristics. Let us introduce the two coupled ordinary differential equations

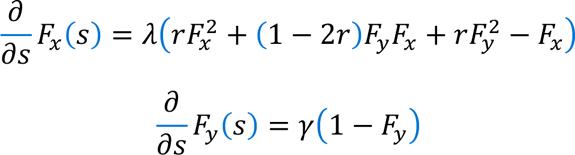

with initial condition *F*_*x*_(0) = *x* and *F*_*y*_(0) = *y*. Then, for each *t* ≥ 0, *G*(*x*, *y*, *t*) = *F*_*x*_(*t*), meaning that the PDE defining *G* is solved using the coupled ODEs defining *F*_*x*_ and *F*_*y*_. Let *p*_*b*_(*t*) = ∑_*n*+*m*0*b*_ *p*_*nm*_(*t*) be the probability of having a clone with *b* basal layer cells, i.e., *b* progenitor cells and differentiated cells. The probabilities *p*_*b*_(*t*) can be extracted from *G* using complex analysis by defining

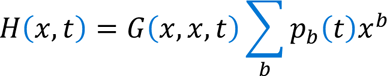

For each fixed *t* > 0, *H*(·, *t*) is a Z-transform of the discrete time probability signal containing all *p*_*b*_(*t*). We extract these probabilities via the inverse Z-transform

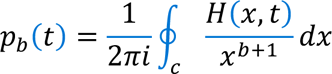

where *c* is an anticlockwise contour contained in the region of convergence and encircling the origin. Since *H*(*x*, *t*) is convergent on the unit disc |*x*| ≤ 1, we integrate over the unit circle, leading to

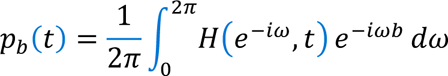

This integral can be approximated using the sum

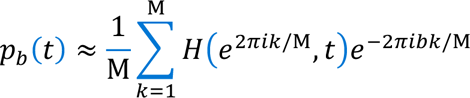

for large enough Μ. This sum can be efficiently computed simultaneously for all relevant clone sizes *b* using a fast Fourier transform of *H*(*e*^2*πik*/M^, *t*), leading to an efficient procedure to find the probablities *p*_*b*_(*t*). The model probabilities in the histograms are found via this approach.

##### Parameter estimation and contour plots

Two different sets of parameters (*r*=0.1, λ=1.9, γ=3.5) ^7^ and (*r*=0.1, λ=2.9, γ=5.5) ^6^ are described to fit the single-progenitor model well. Implementing both sets of parameters renders slightly different average basal clone sizes, as demonstrated in Figure 3E. To estimate the proliferation and stratification parameters in the Troy^WT/KI^ mouse line, we use the more recently published parameters generating larger basal clones (λ=2.9, γ=5.5) as a representation of the Sox2^WT/KI^ data, allowing us to conservatively estimate the increase in *r* required to explain *Troy*-derived basal clones.

Our contour plots are found via the Bayesian estimation also used in ^7^. Let *D* be the observed clonal fate data, then the probability that the model with parameters *λ*, *r*, and *γ* explains the data satisfies

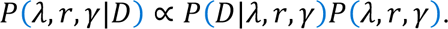

We use a uniform prior *P*(*λ*, *r*, *γ*) over the ranges we consider (implying the prior does not contribute to the estimate) and the likelihood function *P*(*D*|*λ*, *r*, *γ*) comes from a multinomial distribution with a countable number of outcomes

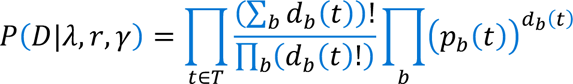

where *d*_*b*_(*t*) is the number of observed clones of size *b* at time *t*, all contained in *D*, *p*_*b*_(*t*) are the clone size probabilities computed as described above, and *T* is a (sub)set of time points for which we have data. Since the data is the same over all estimates for a given *T*, the log- likelihood satisfies

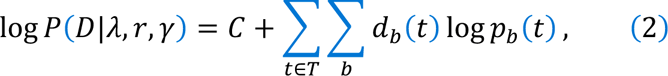

for some constant *C*. After normalization of *d*_*b*_(*t*) to form a probability distribution denoted 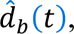 this can equivalently be written as

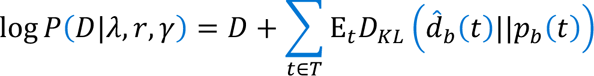

for some constants *D* and *E*, where *D*_*KL*_ denotes the Kullback–Leibler divergence between probability distributions 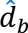 and *p*_*b*_. The contour plots show the sum in (2) for different choices of *T* and parameters *λ*, *r*, and *γ*.

##### Average clone size plots

For the average clone size plots, we simulate (1) forward in time. We run 50 batches, each consisting of 2.000 cell simulations. The average clone size plots show the averages along with standard deviations of the averages in these batches. We compare this to the average clone sizes in different mice, in which 2.000 to 5.000 cells have been counted.

#### RT-qPCR

Tissue samples were collected in Trizol LS Reagent (Thermo Fisher). RNA was extracted using RNAeasy (Qiagen) or Direct-zol (Zymo) and treated with DNaseI (Qiagen). After the quality check, 100ng of total RNA was used for cDNA synthesis using SuperScript IV Vilo (Thermo Fisher). RT-qPCR was run with selected primer pairs and SYBR Green on a ViiA 7 device (Applied biosystems). HPRT and 18sRNA were used as internal controls. Organoids were resuspended in cold PBS and washed 3 times for 5min, lysed in RLT buffer using vortexing to disrupt organoids. RNA was isolated using the RNAeasy mini kit (QIAGEN).

#### RNAScope

RNA in situ hybridization to probe *Tnfrsf19/Troy* and *Axin2* was conducted using a Multiplex Fluorescent V2 kit from Advanced Cell Diagnostics (ACD), according to the manufacturer’s instruction. In brief, fixed frozen sections were allowed to equilibrate to room temperature and washed 2 times for 5 min in PBS. Afterwards, hydrogen peroxide was applied for 10 min at room temperature. The samples were then treated with protease III at 40°C in the oven for 30min, followed by incubation with probes for 2 hours at 40°C in the oven. After amplification with AMP 1, AMP 2, and AMP 3 each channel was developed by incubating HRP, fluorophore, and HRP blocker sequentially based on the channels of the probe that was incubated. Slides were counter-stained with DAPI or immunofluorescent staining as described before. Images were acquired with a Zeiss AxioplanZ2 microscope.

### QUANTIFICATION AND STATISTICAL ANALYSES

All statistical analysis was performed using Prism software (v9.4.1). The specific statistical tests used are indicated in the figure legends along the number of n. Image analysis was performed using Fiji ^50^. Data is represented as mean + SD unless otherwise stated.

### KEY RESOURCE TABLE

| REAGENT or RESOURCE | SOURCE | IDENTIFIER |
| --- | --- | --- |
| Antibodies |  |  |
| E-cadherin (CD324) Antibody | Cell Signaling Technology | Cat #3195 |
| Keratin 5 Polyclonal Chicken Antibody, Purified | BioLegend | Cat #905901 |
| Anti-Cytokeratin 4 antibody [6B10] | abcam | Cat #9004 |
| Anti-Cytokeratin 13 antibody [EPR3671] | abcam | Cat #92551 |
| Anti-SOX2 antibody [EPR3131] | abcam | Cat #92494 |
| Anti-p63 Antibody | abcam | Cat #ab735 |
| Recombinant Anti-KLF4 Antibody | abcam | Cat #ab214666 |
| Anti-Cyclin A2 antibody [EPR17351] | abcam | Cat #181591 |
| Ki-67 Monoclonal Antibody (SolA15) | eBioscience™ | Cat #14-5698 |
| CD49f (Integrin alpha 6) Monoclonal Antibody (eBioGoH3 (GoH3)) | eBioscience™ | Cat #14-0495 |
| Alexa Fluor® 647 anti-mouse CD104 Antibody | BioLegend | Cat #123608 |
| PE/Cy7 anti-mouse/human CD324 (E-Cadherin) Antibody | BioLegend | Cat #147310 |
| Green Fluorescent Protein (GFP) Antibody | Aves Labs, Inc. | Cat # GFP-1020 |
| Chemicals, peptides, and recombinant proteins |  |  |
| SYTOX™ Blue Dead Cell Stain, for flow cytometry | ThermoFisher Scientific | Cat #S34857 |
| Formaldehyde, 37%, with 10-15% methanol | SigmaAldrich | Cat #252549-1L |
| Thermolysin | SigmaAldrich | Cat #T7902-25MG |
| Liberase™ | SigmaAldrich | Cat #5401127001 |
| Noggin murine | Peprtech | Cat #250-38 |
| Rspondin-1 murine | Peprtech | Cat #315-32 |
| Recombinant Mouse EGF Protein, CF | R&D Systems | Cat #2028-EG-200 |
| N-Acetyl-cysteine | SigmaAldrich | Cat # A9165-25G |
| Y-27632 dihydrochloride | SigmaAldrich | Cat # Y0503-5MG |
| Retinoic acid, ≥98% (HPLC), powder | SigmaAldrich | Cat # R2625-100MG |
| Triton™ X-100 solution | SigmaAldrich | Cat # 93443-100ML |
| Tween 20, 100% Nonionic Detergent | Bio-Rad | Cat #1706531 |
| B-27™ Supplement (50X), serum free | ThermoFisher | Cat # 7504001 |
| GlutaMAX™ Supplement | ThermoFisher | Cat #35050061 |
| Penicillin-Streptomycin (10,000 U/mL) | ThermoFisher | Cat #15140122 |
| Dimethyl sulfoxide | SigmaAldrich | Cat #276855-100ML |
| Tamoxifen | SigmaAldrich | Cat #T5648-5G |
| 5-Ethynyl-2'-deoxyuridine | SigmaAldrich | Cat # 900584-50MG |
| Corning® Matrigel® Growth Factor Reduced (GFR) Basement Membrane Matrix | Fisher Scientific™ | Cat #356231 |
| ProLong™ Gold Antifade Mountant 10mL | ThermoFisher | Cat #P36930 |
| DMEM/F-12 | ThermoFisher | Cat #11320074 |
| EDTA | VWR | Cat #E522-100ML |
| HBSS, no calcium, no magnesium, no phenol red | ThermoFisher | Cat #14175-129 |
| DAPI (4',6-Diamidino-2-Phenylindole, Dihydrochloride) | ThermoFisher | Cat # D1306 |
| Critical commercial assays |  |  |
| RNAscope® Multiplex Fluorescent Detection Kit v2 | ACD | Cat # 323110 |
| RNAscope® Probe- Mm-Tnfrsf19 | ACD | Cat # 420241 |
| RNAscope® Probe- Mm-Axin2 | ACD | Cat # 400331-C2 |
| Deposited data |  |  |
| Single-cell sequencing | GEO | GSE255402 |
| RNA-sequencing | GEO | GSE255401 |
| Experimental models: Organisms/strains |  |  |
| TroyCreERT2;tdTomato | Hans Clevers, Hubrecht Institute | PMID: 24120136 |
| Confetti reporter | Jackson Laboratory | JAX:013731 |
| B6;129S-Sox2tm1(cre/ERT2)Hoch/J | Jackson Laboratory | JAX:017593 |
| B6.129S4-*Pdgfra*<sup>tm11(EGFP)Sor</sup>/J | Jackson Laboratory | JAX:007669 |
| Software and algorithms |  |  |
| Prism (v9.4.1) | GraphPad Software | N/A |
| Fiji (v1.0) | ImageJ | N/A |
| Code for cell fate modelling | GitHub | https://doi.org/10.5281/zenodo.10781132 |

## Supplemental Table Legends

**Supplemental Table 1: Differentially expressed genes in single-cell clusters 1-7. Related to Figure 2**.

List of differentially expressed genes identified with RaceID between esophageal epithelial cell clusters comprised of gene name, mean expression in cells not included in cluster, mean expression in cells within cluster, fold-change, p-value, adjusted p-value, and cluster. All genes with a p-value < 0.05 are included.

**Supplemental Table 2: GO terms for genes enriched in cluster 1. Related to Figure 2**.

List of enriched biological process GO terms identified based on differentially expressed genes in cluster 1 compared to cluster 2-7. All GO terms with an adjusted p-value < 0.05 are included.

**Supplemental Table 3: Differentially expressed genes in Troy+ progenitor cells. Related to Figure 2**.

List of the 54 differentially expressed genes between Troy-positive and Troy-negative cells in cluster 1.

**Supplemental Table 4: GO terms enriched in Troy+ cells in cluster 1. Related to Figure 2**.

List of enriched biological process GO terms identified based on differentially expressed genes between Troy-positive and Troy-negative cells in cluster 1.

**Supplemental Table 5: Differentially expressed genes in Troy^KI/KI^ progenitor cells. Related to Figure 4**.

List of differentially expressed genes identified with DESeq2 between FACS assisted isolation of E-CADHERIN-positive, INTEGRINα6-positive, tdTOMATO reporter-positive cells from Troy^WT/KI^ and Troy^KI/KI^ mice.

**Supplemental Table 6: GO terms associated with Troy^KI/KI^ gene expression. Related to Figure 4**.

List of enriched biological process (BP), cellular compartment (CC), and molecular function (MF) GO terms identified based on differentially expressed genes between tdTOMATO reporter-positive cells of Troy^WT/KI^ and Troy^KI/KI^ mice.

## Supplemental Figure legends

**Figure S1:**
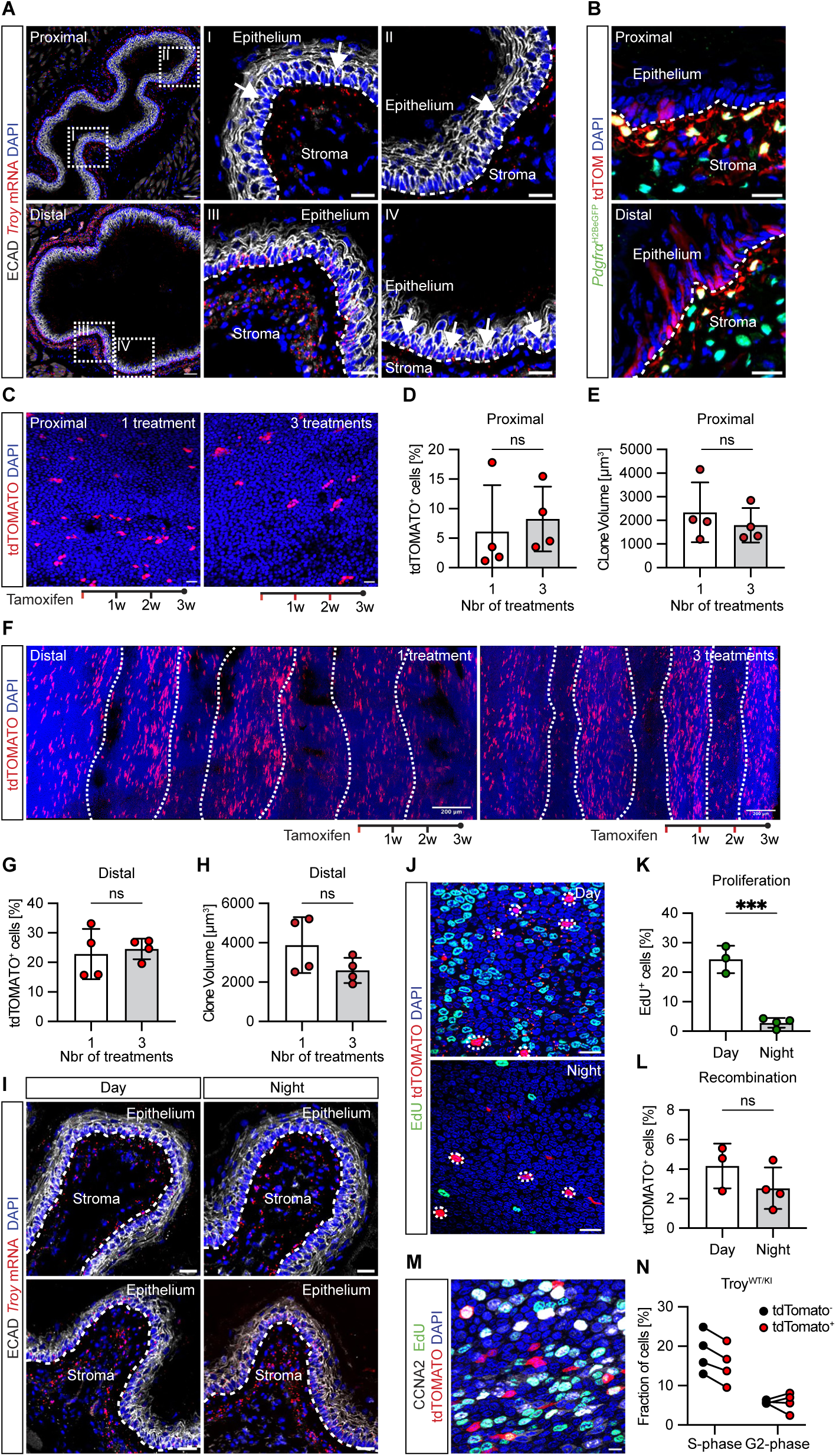
*Troy* delineates a regionally restricted population of basal cells. Related to Figure 1. **(A)** *Troy* mRNA localization in the proximal and distal esophageal epithelium. Arrows point to *Troy* expressing cells. **(B)** Co-expression of tdTOMATO reporter expression and *Pdgfr*α:H2BGFP-positive stromal cells, indicating that a subset of fibroblasts express *Troy*. **(C-H)** Reporter expression comparing one or three tamoxifen injections (once weekly) in Troy^WT/KI^ esophagi. **(C)** Whole mount preparation of the proximal esophagus comparing one and three tamoxifen injections. **(D-E)** Quantification of the contribution of tdTOMATO-labelled cells **(D)** or the average clone volume **(E)** in the proximal esophagus after one or three injections. **(F)** Whole mount preparation of the distal esophagus comparing tdTOMATO abundance and expression **(G-H)** after one or three tamoxifen injections. White lines delineate folding of the epithelium. Neither the percentage of tdTOMATO reporter-positive cells, nor the clone volume, is altered in the proximal or distal esophagus when comparing one and three injections. **(I)** *Troy* mRNA localization in the proximal and distal esophageal epithelium during day and night. **(J)** tdTOMATO reporter expression and EdU incorporation in Troy^WT/KI^ esophagi administered during the day or night. (**K)** Quantification of EdU incorporation (2 hours pulse) comparing day and night proliferation in Troy^WT/KI^ esophagi. **(L)** Administration of tamoxifen during night does not alter the percentage of recombined tdTOMATO-positive cells when compared to reporter expression induced at daytime. (**M-N)** Combining a 2-hour EdU pulse (S-phase) with immunofluorescent localization of CYCLINA2 (expressed in G2 and S-phase) allows for the discrimination of the two cell cycle phases. *Troy*-positive progenitor cells are not preferentially located in G2 phase (CYCLINA2 positive, EdU negative cells). Data are represented as mean ± SD. ^∗^p< 0.05, ^∗∗^p<0.01, ^∗∗∗^p < 0.001, ^∗∗∗∗^ p < 0.0001. Statistical tests are: Two-sided unpaired t-test (**D, E, G, H, K, and L**). Scale bars are: 10μm (**M**), 20μm (**B, I** and **J,** as well as inlets in **A**) 50μm (**A and C**), and 200μm (**F**).

**Figure S2:**
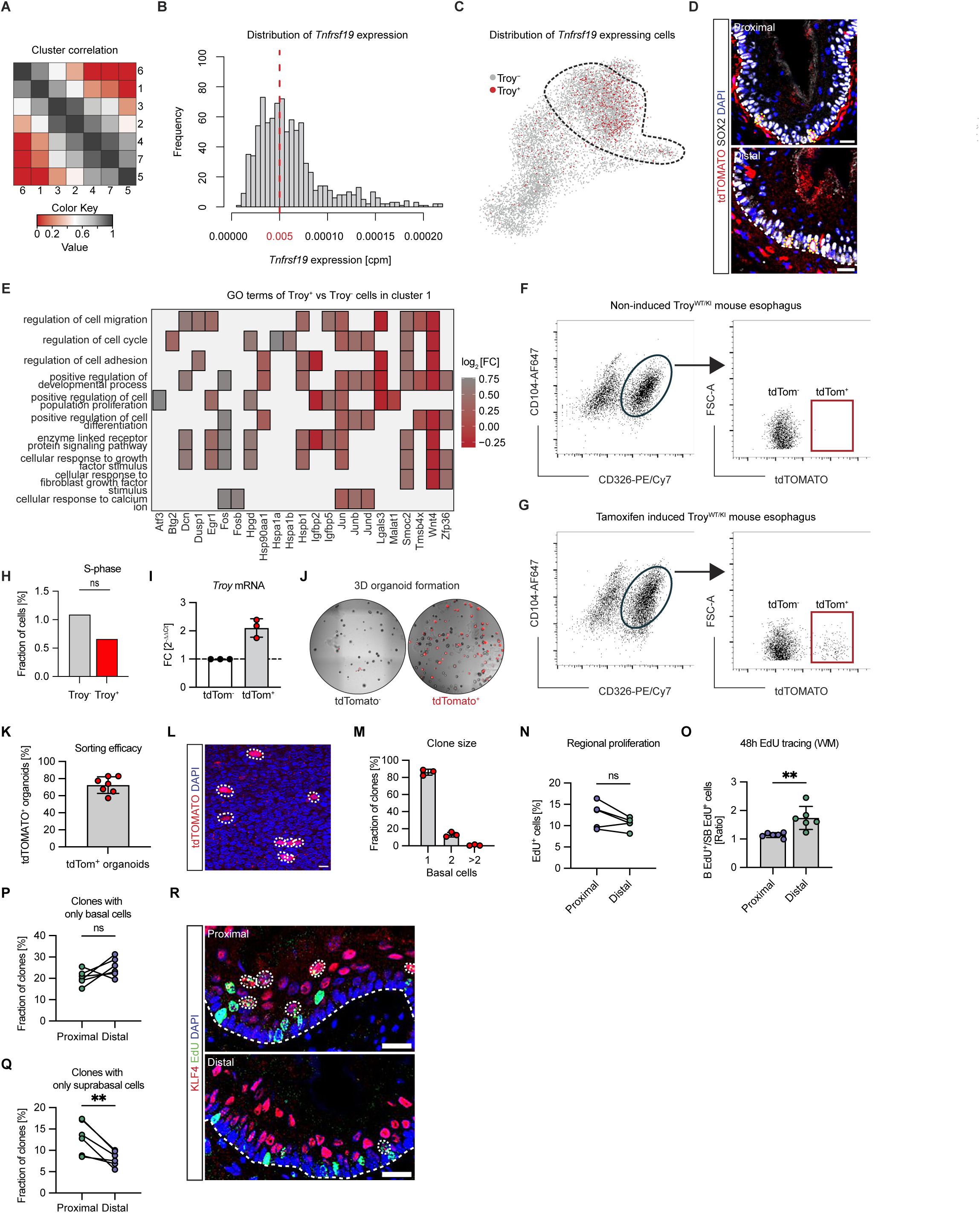
*Troy* marks a functionally distinct progenitor population. Related to Figure 2. **(A)** Cluster correlation indicating the differentiation trajectory of identified clusters. **(B)** *Troy*-positive cells were defined as having >0.005 *Tnfrsr19* counts per million (cpm). **(C)** UMAP visualizing *Troy*-positive (ref) cells. **(D)** *Troy* reporter-positive cells in the basal layer express SOX2. **(E)** Heatmap displaying differentially expressed genes in *Troy*-positive and negative cells from cluster 1, highlighting that many genes are linked to multiple GO terms. **(F)** FACS isolation strategy allowing enrichment of ITGb4/ECAD (CD104/CD326) double-positive basal progenitors, some of which are tdTOMATO-positive after tamoxifen administration **(G)**. **(H)** Analysis of S-phase associated genes in Troy-positive and Troy-negative progenitor cells. **(I)** qPCR analysis of *Troy m*RNA in tdTOMATO-positive and negative FACS isolated progenitor cells. **(J)** Organoids formed from FACS isolated tdTOMATO-positive and negative progenitor cells. **(K)** Percentage of tdTOMATO-positive organoids derived from tdTOMATO sorted progenitor cells. **(L)** Reporter-positive progenitor distribution 24 hours after tamoxifen administration. **(M)** Quantification reveals that most clones are single cells, whereas only a subset consists of 2 or more basal cells. **(N)** Quantification of EdU-positive cells comparing proximal and distal esophagus after a 1-hour EdU pulse reveal no statistically significant difference in proliferation rates. **(O)** Quantification of all EdU-positive daughter cells after 48 hours of EdU tracing comparing proximal and distal esophagus show that distal progenitor cells generate more basal, compared to suprabasal, progeny. **(P-Q)** Clones consisting of only basal cells are not significantly different comparing proximal and distal esophagus at 48 hours of EdU tracing. However, clones consisting of suprabasal cells only are enriched in the proximal esophagus, indicating that progenitor cell behavior in the proximal and distal esophagus is distinct. **(R)** Representative image of clone distribution in proximal and distal esophagus after 48 hours or EdU tracing. Data are represented as mean ± SD. ^∗^p< 0.05, ^∗∗^p<0.01, ^∗∗∗^p < 0.001, ^∗∗∗∗^ p < 0.0001. Statistical tests are: Wilcoxon rank sum test (**H**), Wilcoxon matched-pairs signed-ranked test (**N-Q**). Scale bars are: 20μm (**D, L** and **R**).

**Figure S3.**
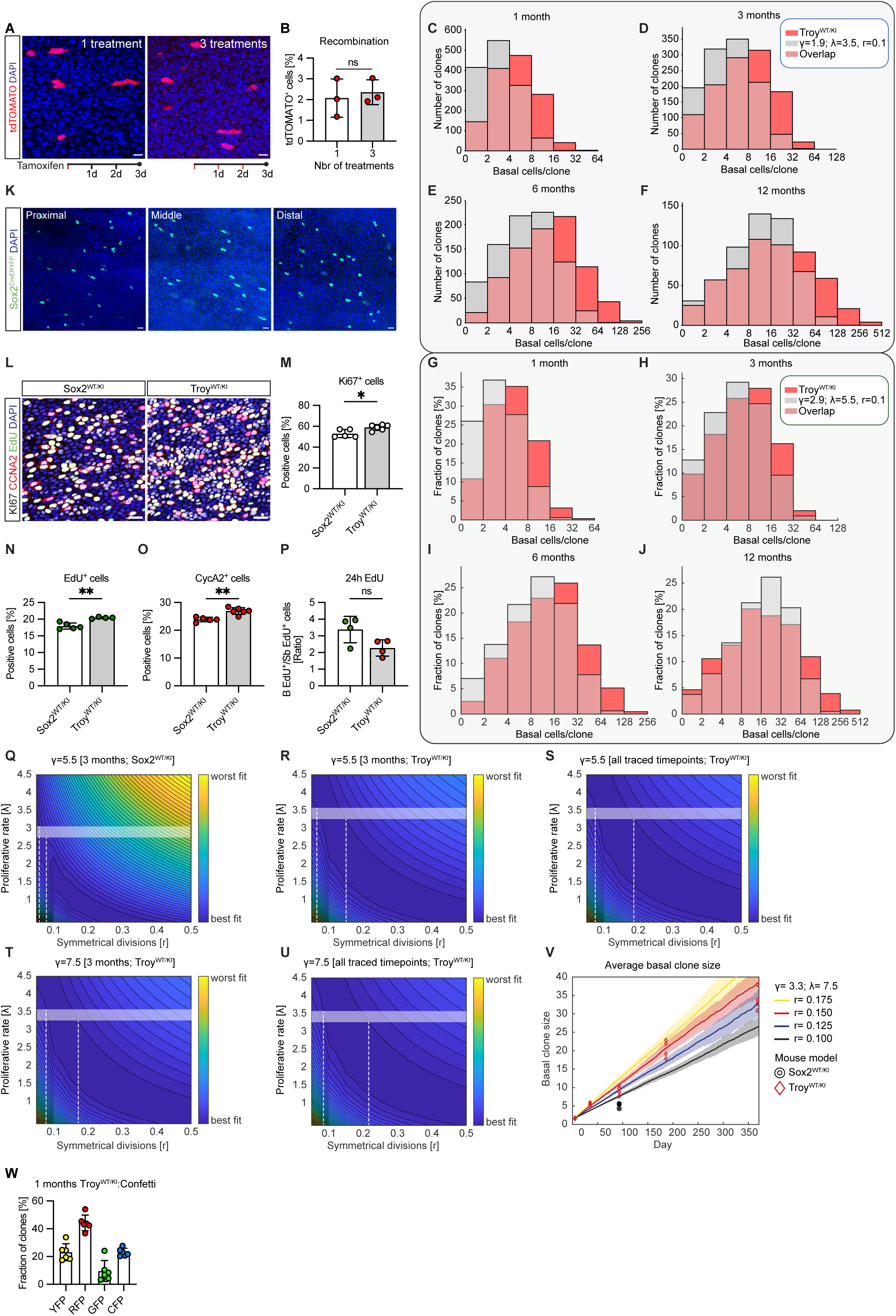
*Troy*-positive progenitors contribute to tissue homeostasis long-term. Related to Figure 3. **(A-B)** Topical tamoxifen treatment for one or three consecutive days does not significantly alter the number of labelled basal cells. **(C-F)** Histograms displaying the number of clones of specific basal cell sizes at 1, 3, 6 and 12 months. Red bars represent the experimental data derived from *Troy-CreER^T2^:tdTomato* (Troy^WT/KI^) mice, whereas grey bars indicate the predicted clone size distributions from the single-progenitor, or nominal cell model (λ=1.9 : γ=3.5). **(G-J)** Histograms displaying the number of clones of specific basal cell sizes at 1, 3, 6 and 12 months. Red bars represent the experimental data derived from *Troy-CreER^T2^:tdTomato* (Troy^WT/KI^) mice, whereas grey bars indicate the predicted clone size distributions from the single-progenitor, or nominal cell model (λ=2.9 : γ=5.5). **(C-J)** *Troy*-derived clones are larger than would be predicted at all time points. **(K)** Topical administration of tamoxifen to *Sox2-CreER^T2^:YFP* (Sox2^WT/KI^) reveals even distribution of recombination along the esophageal axis after 24 hours chase. **(L-O)** Representative images and quantification comparing KI67, CYCLINA2 (CCNA2) and EdU in Sox2^WT/KI^ and Troy^WT/KI^ mice. **(P)** Quantification of EdU-positive cells (basal and suprabasal) after 24 hours of chase in Sox2^WT/KI^ and Troy^WT/KI^ mice. **(Q)** Contour plot visualizing the distribution of optimal parameters at 3 months of tracing in the Sox2^WT/KI^ mouse (γ=5.5). Best fit probabilities for *r* are dark blue and correlate to (r <0.1) within the experimentally determined proliferation span (λ=2.9). Horizontally shaded area marks the experimentally predicted proliferation rate, dotted white lines highlight corresponding optimal *r*-probabilities. **(R-U)** Contour plots visualizing the distribution of optimal parameters at in the Troy^WT/KI^ mouse using (γ=5.5) (**R** and **S**) or (γ=7.5) (**T** and **U**). **(V)** Average basal clone size prediction modelling increasing r (r=0.1-0.175) with fixed proliferation and stratification rates (λ=3.3 : γ=7.5). **(W)** Distribution of reporter positive (YFP, RFP, GFP and CFP) basal clones after 1 month of lineage tracing in the Confetti reporter line. Data are represented as mean ± SD. ^∗^p< 0.05, ^∗∗^p<0.01, ^∗∗∗^p < 0.001, ^∗∗∗∗^ p < 0.0001. Shaded areas in (**V**) indicate SD based on simulations. Statistical tests are: Two-sided unpaired t-test (**B, M, N, O, P**). Scale bars are: 20μm (**A** and **L**) and 50μm (**K**). n=3 mice in **B**, n=3-6 mice in **C-J, M-P, V and W**.

**Figure S4.**
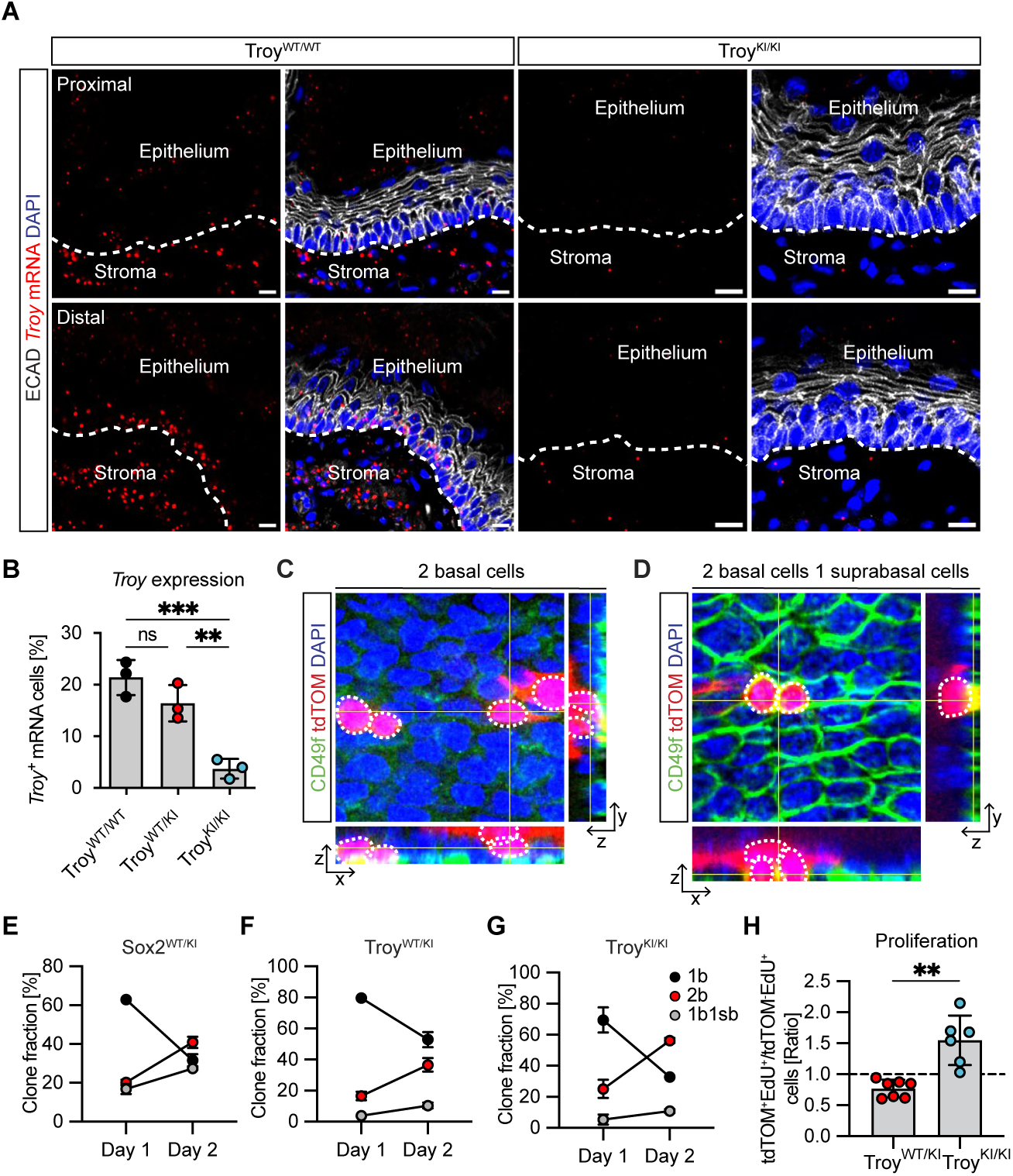
*Troy* negatively regulates progenitor cell proliferation. Related to Figure 4. **(A)** Visualization of *Troy* mRNA in Troy^WT/KI^ and Troy^KI/KI^ proximal and distal esophagus. **(B)** Comparison of Troy mRNA levels in Troy^WT/WT^, Troy^WT/KI^ and Troy^KI/KI^. Troy expression is retained in Troy^WT/KI^ compared to Troy^WT/WT^, however significantly reduced in Troy^KI/KI^ when compared to either Troy^WT/WT^ or Troy^WT/KI^. **(C-D)** Representative 3D visualization of tdTOMATO-positive 2-cell basal only **(C)** or 3-cell (2 basal + 1 suprabasal cell) clone **(D)**. CD49f is used to delineate the basal layer. **(E-G)** Clonal development comparing 1 and 2 days of lineage tracing in Sox2^WT/KI^, Troy^WT/KI^and Troy^KI/KI^ mouse lines, focusing on fate changes linked to the first division. Whereas all three lines display a reduction in clones consisting of 1 basal cell (1b) and an increase in two cell (2b or 1b1sb) clones, the dynamics are distinct. Sox2^WT/KI^ display an increased fraction of asymmetrically fated (1b1sb, grey circles) clones compared to Troy^WT/KI^ and Troy^KI/KI^ -derived clones. **(H)** Relative proliferation profiles in tdTOMATO reporter-positive progenitors in Troy^WT/KI^ and Troy^KI/KI^ esophagi, compared to their respective tdTOMATO-negative progenitor population. Data are represented as mean ± SD. ^∗^p< 0.05, ^∗∗^p<0.01, ^∗∗∗^p < 0.001, ^∗∗∗∗^ p < 0.0001. Statistical tests are: Ordinary one-way ANOVA (**B**), Two-sided unpaired t-test (**H**). Scale bars are: 10μm (**C and D**), 20μm (**A**). n=3-5 mice in **D-F.**

**Figure S5.**
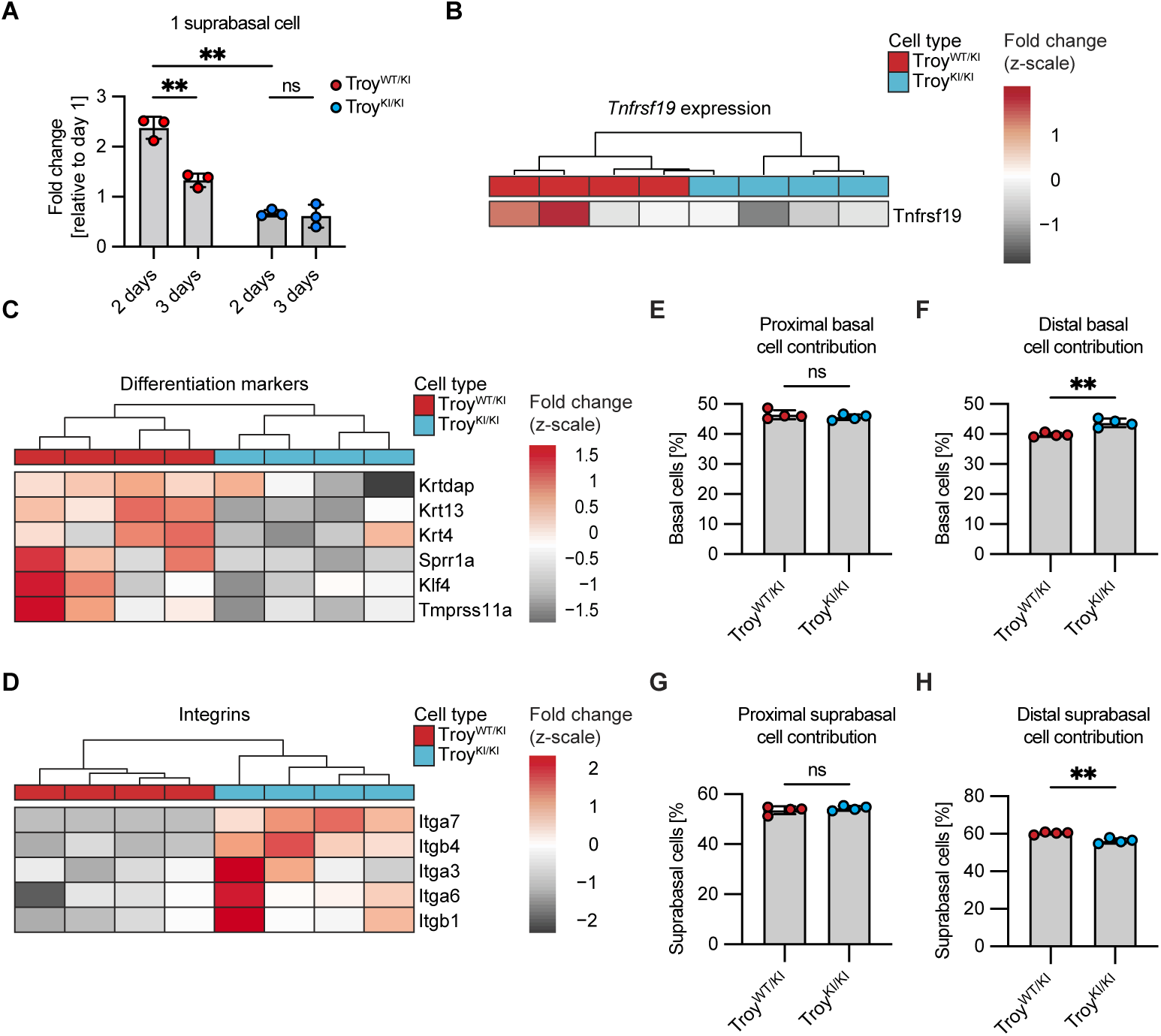
Commitment to differentiation is impaired in the absence of *Troy.* Related to Figure 5. **(A)** Behavior of single suprabasal cell clones at 2 and 3 days compared to day 1. Troy^WT/KI^ cells are more prone to delaminate when compared to Troy^KI/KI^ cells. **(B)** Bulk RNA-sequencing data confirm loss of Troy expression in Troy^KI/KI^ compared to Troy^WT/KI^ tdTOMOATO cells. **(C)** Differentiation markers are reduced in the absence of TROY. **(D)** Expression of integrins is increased in Troy^KI/KI^ when compared to Troy^WT/KI^ progenitor cells. **(E-F)** Percentage of basal cells (out of all cells) in proximal and distal esophagus, comparing Troy^WT/KI^ and Troy^KI/KI^, indicating that the Troy^KI/KI^ basal cell population is expanded. **(G-H)** Percentage of suprabasal cells (out of all cells) in proximal and distal esophagus, comparing Troy^WT/KI^ and Troy^KI/KI^, indicating that the Troy^KI/KI^ suprabasal cell population is reduced. Data are represented as mean ± SD. ^∗^p< 0.05, ^∗∗^p<0.01, ^∗∗∗^p < 0.001, ^∗∗∗∗^ p < 0.0001. Statistical tests are: Multiple unpaired t-tests (adjusted p-value < 0.05) (**A**), Two-sided unpaired t-test (**E-H**). n=3-5 mice in **E-H.**

**Figure S6.**
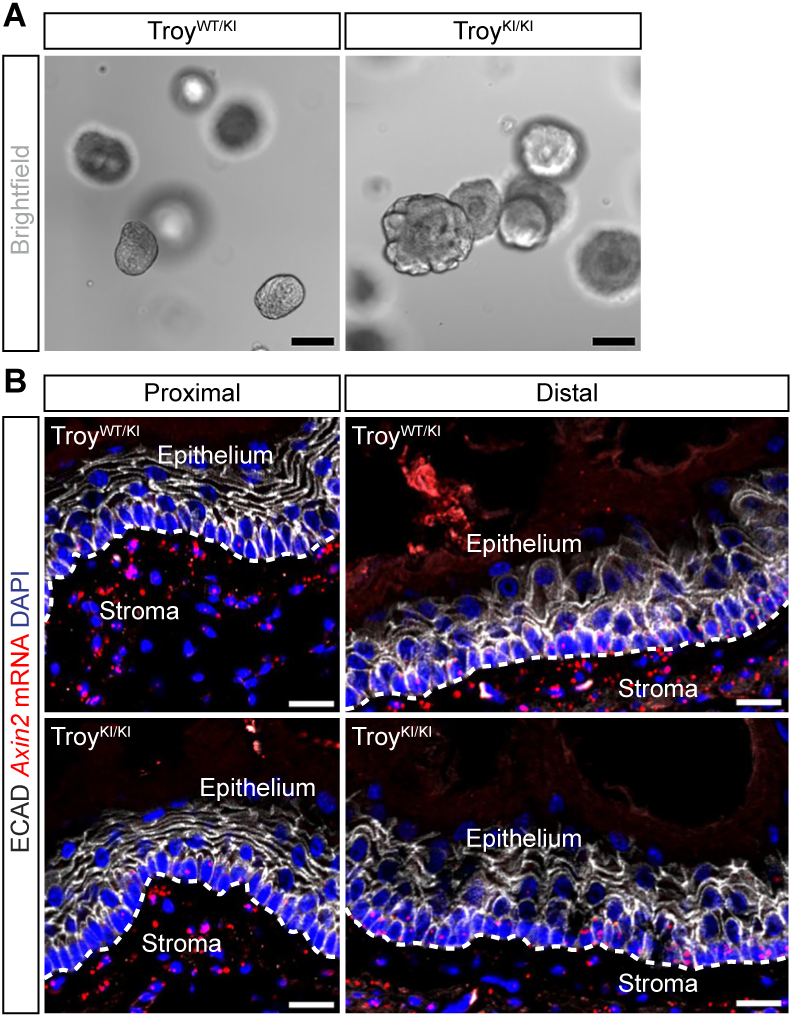
Troy^KI/KI^ proliferation dynamics are cell autonomous. Related to Figure 6. **(A)** Brightfield representation of organoids derived from Troy^WT/KI^ and Troy^KI/KI^. **(B)** Visualization of *Axin2* mRNA in proximal and distal esophagus. Loss of TROY does not affect *Axin2* expression. Scale bars are: 20μm (**B**) and 200μm (**A**).

**Figure S7:**
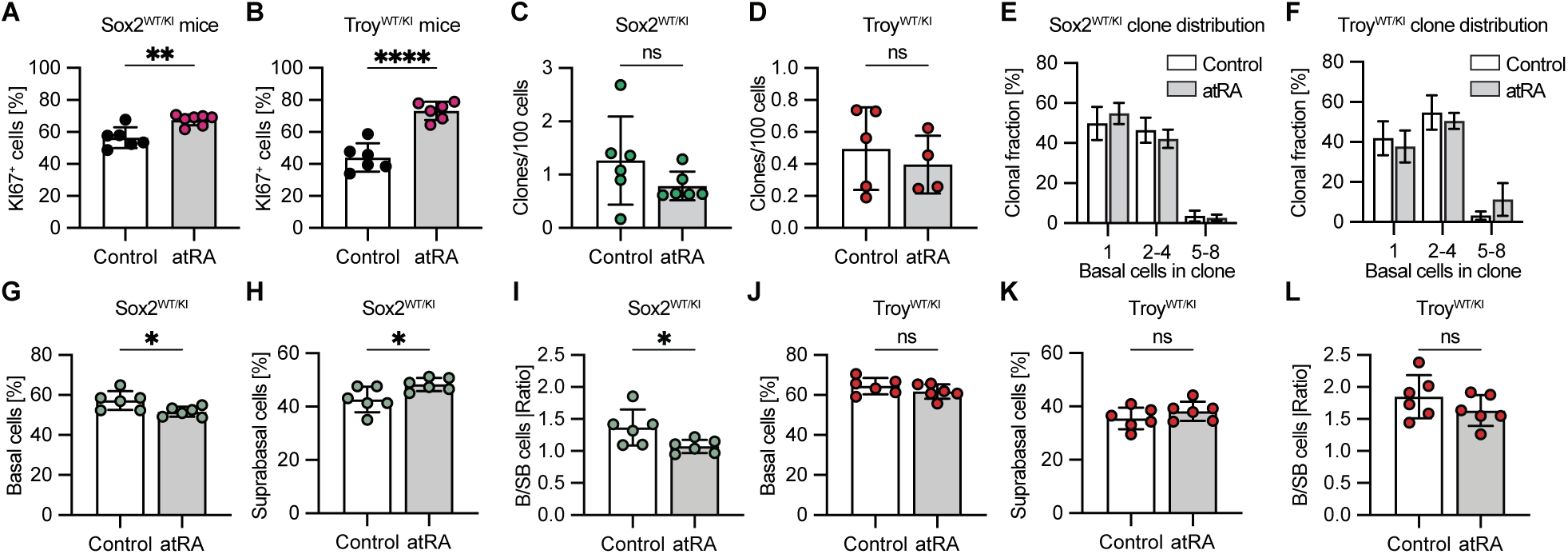
*Troy* is dynamically regulated upon tissue stress. Related to Figure 7. **(A-B)** Quantification of KI67-positive cells in Sox2^WT/KI^ and Troy^WT/KI^ esophagi confirm that atRA treatment induce hyperproliferation in the esophageal epithelium when compared to control treated esophagi. **(C-D)** atRA treatment does not alter the number of remaining clones at day 9, suggesting that *Troy* expression is silenced during atRA (see Figure 4C). **(E-F)** Clone size distribution in Sox2^WT/KI^ and Troy^WT/KI^ is not significantly altered when comparing control to atRA treated esophagi within the same line. **(G-I)** Quantification of the basal **(G)** and suprabasal **(H)** cell contribution to Sox2^WT/KI^ clones, comparing control and atRA treatment. atRA shifts Sox2^WT/KI^-derived clones towards differentiation (**I**). **(J-K)** Quantification of the basal (**J**) and suprabasal (**K**) cell contribution to Troy^WT/KI^ clones, comparing control and atRA treatment. Troy^WT/KI^-derived clones are refractory to atRA-induced differentiation (**L**). Data are represented as mean ± SD. ^∗^p< 0.05, ^∗∗^p<0.01, ^∗∗∗^p < 0.001, ^∗∗∗∗^ p < 0.0001. Statistical tests are: Two-sided unpaired t-test (**A-D, G-L**). n=6 mice.

## References

1. Croagh, D., Phillips, W.A., Redvers, R., Thomas, R.J., and Kaur, P. (2007). Identification of candidate murine esophageal stem cells using a combination of cell kinetic studies and cell surface markers. Stem Cells 25, 313–318. 10.1634/stemcells.2006-0421.

2. DeWard, A.D., Cramer, J., and Lagasse, E. (2014). Cellular heterogeneity in the mouse esophagus implicates the presence of a nonquiescent epithelial stem cell population. Cell Rep 9, 701–711. 10.1016/j.celrep.2014.09.027.

3. Giroux, V., Lento, A.A., Islam, M., Pitarresi, J.R., Kharbanda, A., Hamilton, K.E., Whelan, K.A., Long, A., Rhoades, B., Tang, Q., et al. (2017). Long-lived keratin 15+ esophageal progenitor cells contribute to homeostasis and regeneration. J Clin Invest 127, 2378–2391. 10.1172/JCI88941.

4. Kalabis, J., Oyama, K., Okawa, T., Nakagawa, H., Michaylira, C.Z., Stairs, D.B., Figueiredo, J.L., Mahmood, U., Diehl, J.A., Herlyn, M., and Rustgi, A.K. (2008). A subpopulation of mouse esophageal basal cells has properties of stem cells with the capacity for self-renewal and lineage specification. J Clin Invest 118, 3860–3869. 10.1172/JCI35012.

5. Jones, K.B., Furukawa, S., Marangoni, P., Ma, H., Pinkard, H., D’Urso, R., Zilionis, R., Klein, A.M., and Klein, O.D. (2019). Quantitative Clonal Analysis and Single-Cell Transcriptomics Reveal Division Kinetics, Hierarchy, and Fate of Oral Epithelial Progenitor Cells. Cell Stem Cell 24, 183–192 e188. 10.1016/j.stem.2018.10.015.

6. Piedrafita, G., Kostiou, V., Wabik, A., Colom, B., Fernandez-Antoran, D., Herms, A., Murai, K., Hall, B.A., and Jones, P.H. (2020). A single-progenitor model as the unifying paradigm of epidermal and esophageal epithelial maintenance in mice. Nat Commun 11, 1429. 10.1038/s41467-020-15258-0.

7. Doupe, D.P., Alcolea, M.P., Roshan, A., Zhang, G., Klein, A.M., Simons, B.D., and Jones, P.H. (2012). A single progenitor population switches behavior to maintain and repair esophageal epithelium. Science 337, 1091–1093. 10.1126/science.1218835.

8. Cockburn, K., Annusver, K., Gonzalez, D.G., Ganesan, S., May, D.P., Mesa, K.R., Kawaguchi, K., Kasper, M., and Greco, V. (2022). Gradual differentiation uncoupled from cell cycle exit generates heterogeneity in the epidermal stem cell layer. Nat Cell Biol 24, 1692–1700. 10.1038/s41556-022-01021-8.

9. Mascre, G., Dekoninck, S., Drogat, B., Youssef, K.K., Brohee, S., Sotiropoulou, P.A., Simons, B.D., and Blanpain, C. (2012). Distinct contribution of stem and progenitor cells to epidermal maintenance. Nature 489, 257–262. 10.1038/nature11393.

10. Barker, N., Ridgway, R.A., van Es, J.H., van de Wetering, M., Begthel, H., van den Born, M., Danenberg, E., Clarke, A.R., Sansom, O.J., and Clevers, H. (2009). Crypt stem cells as the cells-of-origin of intestinal cancer. Nature 457, 608–611. 10.1038/nature07602.

11. Lapouge, G., Youssef, K.K., Vokaer, B., Achouri, Y., Michaux, C., Sotiropoulou, P.A., and Blanpain, C. (2011). Identifying the cellular origin of squamous skin tumors. Proc Natl Acad Sci U S A 108, 7431–7436. 10.1073/pnas.1012720108.

12. Jiang, M., Ku, W.Y., Zhou, Z., Dellon, E.S., Falk, G.W., Nakagawa, H., Wang, M.L., Liu, K., Wang, J., Katzka, D.A., et al. (2015). BMP-driven NRF2 activation in esophageal basal cell differentiation and eosinophilic esophagitis. J Clin Invest 125, 1557–1568. 10.1172/JCI78850.

13. Ohashi, S., Natsuizaka, M., Yashiro-Ohtani, Y., Kalman, R.A., Nakagawa, M., Wu, L., Klein-Szanto, A.J., Herlyn, M., Diehl, J.A., Katz, J.P., et al. (2010). NOTCH1 and NOTCH3 coordinate esophageal squamous differentiation through a CSL-dependent transcriptional network. Gastroenterology 139, 2113–2123. 10.1053/j.gastro.2010.08.040.

14. McGinn, J., Hallou, A., Han, S., Krizic, K., Ulyanchenko, S., Iglesias-Bartolome, R., England, F.J., Verstreken, C., Chalut, K.J., Jensen, K.B., et al. (2021). A biomechanical switch regulates the transition towards homeostasis in oesophageal epithelium. Nat Cell Biol 23, 511–525. 10.1038/s41556-021-00679-w.

15. Zhang, Y., Jiang, M., Kim, E., Lin, S., Liu, K., Lan, X., and Que, J. (2017). Development and stem cells of the esophagus. Semin Cell Dev Biol 66, 25–35. 10.1016/j.semcdb.2016.12.008.

16. Fafilek, B., Krausova, M., Vojtechova, M., Pospichalova, V., Tumova, L., Sloncova, E., Huranova, M., Stancikova, J., Hlavata, A., Svec, J., et al. (2013). Troy, a tumor necrosis factor receptor family member, interacts with lgr5 to inhibit wnt signaling in intestinal stem cells. Gastroenterology 144, 381–391. 10.1053/j.gastro.2012.10.048.

17. Schutgens, F., Rookmaaker, M.B., Blokzijl, F., van Boxtel, R., Vries, R., Cuppen, E., Verhaar, M.C., and Clevers, H. (2017). Troy/TNFRSF19 marks epithelial progenitor cells during mouse kidney development that continue to contribute to turnover in adult kidney. Proc Natl Acad Sci U S A 114, E11190–E11198. 10.1073/pnas.1714145115.

18. Stange, D.E., Koo, B.K., Huch, M., Sibbel, G., Basak, O., Lyubimova, A., Kujala, P., Bartfeld, S., Koster, J., Geahlen, J.H., et al. (2013). Differentiated Troy+ chief cells act as reserve stem cells to generate all lineages of the stomach epithelium. Cell 155, 357–368. 10.1016/j.cell.2013.09.008.

19. Kretzschmar, K., Boonekamp, K.E., Bleijs, M., Asra, P., Koomen, M., Chuva de Sousa Lopes, S.M., Giovannone, B., and Clevers, H. (2021). Troy/Tnfrsf19 marks epidermal cells that govern interfollicular epidermal renewal and cornification. Stem Cell Reports 16, 2379–2394. 10.1016/j.stemcr.2021.07.007.

20. Stenudd, M., Sabelstrom, H., Llorens-Bobadilla, E., Zamboni, M., Blom, H., Brismar, H., Zhang, S., Basak, O., Clevers, H., Goritz, C., et al. (2022). Identification of a discrete subpopulation of spinal cord ependymal cells with neural stem cell properties. Cell Rep 38, 110440. 10.1016/j.celrep.2022.110440.

21. Basak, O., Krieger, T.G., Muraro, M.J., Wiebrands, K., Stange, D.E., Frias-Aldeguer, J., Rivron, N.C., van de Wetering, M., van Es, J.H., van Oudenaarden, A., et al. (2018). Troy+ brain stem cells cycle through quiescence and regulate their number by sensing niche occupancy. Proc Natl Acad Sci U S A 115, E610–E619. 10.1073/pnas.1715911114.

22. Shao, Z., Browning, J.L., Lee, X., Scott, M.L., Shulga-Morskaya, S., Allaire, N., Thill, G., Levesque, M., Sah, D., McCoy, J.M., et al. (2005). TAJ/TROY, an orphan TNF receptor family member, binds Nogo-66 receptor 1 and regulates axonal regeneration. Neuron 45, 353–359. 10.1016/j.neuron.2004.12.050.

23. Deng, C., Lin, Y.X., Qi, X.K., He, G.P., Zhang, Y., Zhang, H.J., Xu, M., Feng, Q.S., Bei, J.X., Zeng, Y.X., and Feng, L. (2018). TNFRSF19 Inhibits TGFbeta Signaling through Interaction with TGFbeta Receptor Type I to Promote Tumorigenesis. Cancer Res 78, 3469–3483. 10.1158/0008-5472.CAN-17-3205.

24. Eby, M.T., Jasmin, A., Kumar, A., Sharma, K., and Chaudhary, P.M. (2000). TAJ, a novel member of the tumor necrosis factor receptor family, activates the c-Jun N-terminal kinase pathway and mediates caspase-independent cell death. J Biol Chem 275, 15336–15342. 10.1074/jbc.275.20.15336.

25. Paulino, V.M., Yang, Z., Kloss, J., Ennis, M.J., Armstrong, B.A., Loftus, J.C., and Tran, N.L. (2010). TROY (TNFRSF19) is overexpressed in advanced glial tumors and promotes glioblastoma cell invasion via Pyk2-Rac1 signaling. Mol Cancer Res 8, 1558–1567. 10.1158/1541-7786.MCR-10-0334.

26. Scheving, L.A. (2000). Biological clocks and the digestive system. Gastroenterology 119, 536–549. 10.1053/gast.2000.9305.

27. Kabir, M.F., Karami, A.L., Cruz-Acuna, R., Klochkova, A., Saxena, R., Mu, A., Murray, M.G., Cruz, J., Fuller, A.D., Clevenger, M.H., et al. (2022). Single cell transcriptomic analysis reveals cellular diversity of murine esophageal epithelium. Nat Commun 13, 2167. 10.1038/s41467-022-29747-x.

28. Grun, D., Lyubimova, A., Kester, L., Wiebrands, K., Basak, O., Sasaki, N., Clevers, H., and van Oudenaarden, A. (2015). Single-cell messenger RNA sequencing reveals rare intestinal cell types. Nature 525, 251–255. 10.1038/nature14966.

29. Shvab, A., Haase, G., Ben-Shmuel, A., Gavert, N., Brabletz, T., Dedhar, S., and Ben-Ze’ev, A. (2016). Induction of the intestinal stem cell signature gene SMOC-2 is required for L1-mediated colon cancer progression. Oncogene 35, 549–557. 10.1038/onc.2015.127.

30. Liu, B.Y., Soloviev, I., Huang, X., Chang, P., Ernst, J.A., Polakis, P., and Sakanaka, C. (2012). Mammary tumor regression elicited by Wnt signaling inhibitor requires IGFBP5. Cancer Res 72, 1568–1578. 10.1158/0008-5472.CAN-11-3668.

31. Verma, B.K., and Kondaiah, P. (2020). Regulation of beta-catenin by IGFBP2 and its cytoplasmic actions in glioma. J Neurooncol 149, 209–217. 10.1007/s11060-020-03596-4.

32. Barrandon, Y., and Green, H. (1987). Three clonal types of keratinocyte with different capacities for multiplication. Proc Natl Acad Sci U S A 84, 2302–2306. 10.1073/pnas.84.8.2302.

33. Abby, E., Dentro, S.C., Hall, M.W.J., Fowler, J.C., Ong, S.H., Sood, R., Herms, A., Piedrafita, G., Abnizova, I., Siebel, C.W., et al. (2023). Notch1 mutations drive clonal expansion in normal esophageal epithelium but impair tumor growth. Nat Genet 55, 232–245. 10.1038/s41588-022-01280-z.

34. Reya, T., and Clevers, H. (2005). Wnt signalling in stem cells and cancer. Nature 434, 843–850. 10.1038/nature03319.

35. Buttitta, L., Tanaka, T.S., Chen, A.E., Ko, M.S., and Fan, C.M. (2003). Microarray analysis of somitogenesis reveals novel targets of different WNT signaling pathways in the somitic mesoderm. Dev Biol 258, 91–104. 10.1016/s0012-1606(03)00116-7.

36. Hao, H.X., Xie, Y., Zhang, Y., Charlat, O., Oster, E., Avello, M., Lei, H., Mickanin, C., Liu, D., Ruffner, H., et al. (2012). ZNRF3 promotes Wnt receptor turnover in an R-spondin-sensitive manner. Nature 485, 195–200. 10.1038/nature11019.

37. Koo, B.K., Spit, M., Jordens, I., Low, T.Y., Stange, D.E., van de Wetering, M., van Es, J.H., Mohammed, S., Heck, A.J., Maurice, M.M., and Clevers, H. (2012). Tumour suppressor RNF43 is a stem-cell E3 ligase that induces endocytosis of Wnt receptors. Nature 488, 665–669. 10.1038/nature11308.

38. Wielenga, V.J., Smits, R., Korinek, V., Smit, L., Kielman, M., Fodde, R., Clevers, H., and Pals, S.T. (1999). Expression of CD44 in Apc and Tcf mutant mice implies regulation by the WNT pathway. Am J Pathol 154, 515–523. 10.1016/S0002-9440(10)65297-2.

39. Chang, C.L., Lao-Sirieix, P., Save, V., De La Cueva Mendez, G., Laskey, R., and Fitzgerald, R.C. (2007). Retinoic acid-induced glandular differentiation of the oesophagus. Gut 56, 906–917. 10.1136/gut.2006.097915.

40. Dekoninck, S., and Blanpain, C. (2019). Stem cell dynamics, migration and plasticity during wound healing. Nat Cell Biol 21, 18–24. 10.1038/s41556-018-0237-6.

41. Snippert, H.J., van der Flier, L.G., Sato, T., van Es, J.H., van den Born, M., Kroon-Veenboer, C., Barker, N., Klein, A.M., van Rheenen, J., Simons, B.D., and Clevers, H. (2010). Intestinal crypt homeostasis results from neutral competition between symmetrically dividing Lgr5 stem cells. Cell 143, 134–144. 10.1016/j.cell.2010.09.016.

42. Arnold, K., Sarkar, A., Yram, M.A., Polo, J.M., Bronson, R., Sengupta, S., Seandel, M., Geijsen, N., and Hochedlinger, K. (2011). Sox2(+) adult stem and progenitor cells are important for tissue regeneration and survival of mice. Cell Stem Cell 9, 317–329. 10.1016/j.stem.2011.09.001.

43. Klinghoffer, R.A., Hamilton, T.G., Hoch, R., and Soriano, P. (2002). An allelic series at the PDGFalphaR locus indicates unequal contributions of distinct signaling pathways during development. Dev Cell 2, 103–113. 10.1016/s1534-5807(01)00103-4.

44. Genander, M., Cook, P.J., Ramskold, D., Keyes, B.E., Mertz, A.F., Sandberg, R., and Fuchs, E. (2014). BMP signaling and its pSMAD1/5 target genes differentially regulate hair follicle stem cell lineages. Cell Stem Cell 15, 619–633. 10.1016/j.stem.2014.09.009.

45. Hsu, Y.C., Pasolli, H.A., and Fuchs, E. (2011). Dynamics between stem cells, niche, and progeny in the hair follicle. Cell 144, 92–105. 10.1016/j.cell.2010.11.049.

46. Vasioukhin, V., Degenstein, L., Wise, B., and Fuchs, E. (1999). The magical touch: genome targeting in epidermal stem cells induced by tamoxifen application to mouse skin. Proc Natl Acad Sci U S A 96, 8551–8556. 10.1073/pnas.96.15.8551.

47. Eenjes, E., Grommisch, D., and Genander, M. (2023). Functional Characterization and Visualization of Esophageal Fibroblasts Using Organoid Co-Cultures. J Vis Exp. 10.3791/64905.

48. Yu, G., Wang, L.G., Han, Y., and He, Q.Y. (2012). clusterProfiler: an R package for comparing biological themes among gene clusters. OMICS 16, 284–287. 10.1089/omi.2011.0118.

49. Picelli, S., Faridani, O.R., Bjorklund, A.K., Winberg, G., Sagasser, S., and Sandberg, R. (2014). Full-length RNA-seq from single cells using Smart-seq2. Nat Protoc 9, 171–181. 10.1038/nprot.2014.006.

50. Schindelin, J., Arganda-Carreras, I., Frise, E., Kaynig, V., Longair, M., Pietzsch, T., Preibisch, S., Rueden, C., Saalfeld, S., Schmid, B., et al. (2012). Fiji: an open-source platform for biological-image analysis. Nat Methods 9, 676–682. 10.1038/nmeth.2019.

